# Higher-order and distributed synergistic functional interactions encode information gain in goal-directed learning

**DOI:** 10.1101/2024.09.23.614484

**Authors:** Etienne Combrisson, Ruggero Basanisi, Matteo Neri, Guillaume Auzias, Giovanni Petri, Daniele Marinazzo, Stefano Panzeri, Andrea Brovelli

**Affiliations:** Institut de Neurosciences de la Timone UMR 7289, Aix Marseille Université, CNRS, 13005, Marseille, France; MoMiLab Research Unit, IMT School for Advanced Studies Lucca, Lucca, Italy; Net/ork Science Institute, Northeastern University, London, London E1W 1LP, United Kingdom; Department of Data Analysis, Faculty of Psychological and Educational Sciences, B-9000 Ghent, Belgium; Institute for Neural Information Processing, University Medical Center Hamburg-Eppendorf (UKE), Hamburg, Germany

**Author notes:** These authors contributed equally (Etienne Combrisson, Ruggero Basanisi).

## Abstract

Goal-directed learning arises from distributed neural circuits including the prefrontal, posterior parietal and temporal cortices. However, the role of cortico-cortical functional interactions remains unclear. To address this question, we integrated information dynamics analysis with magnetoencephalography to investigate the encoding of learning signals through neural interactions. Our findings revealed that information gain (the reduction in uncertainty about the causal relationship between actions and outcomes) is represented over the visual, parietal, lateral prefrontal and ventromedial/orbital prefrontal cortices. Cortico-cortical interactions encoded information gain synergistically at the level of pairwise and higher-order relations, such as triplets and quadruplets. Higher-order synergistic interactions were characterized by long-range relationships centered in the ventromedial and orbitofrontal cortices, which served as key receivers in the broadcast of information gain across cortical circuits. Overall, this study provides evidence that information gain is encoded through synergistic and higher-order functional interactions and is broadcast to prefrontal reward circuits.

## INTRODUCTION

A key factor of human agency is the ability to create beliefs about the consequence of our actions. This ability provides the basis for rational decision-making and, in general, allows people to engage in meaningful life and social interactions all along their life (Bandura, 1997). Learning the causal relation between actions and outcomes is thought to be supported by the goal-directed system (Tolman, 1948; Dickinson, 1994; Dolan and Dayan, 2013). Neurally, goal-directed behaviours emerge from the coordinated activity of neural populations distributed over the associative fronto-striatal circuit (Haber and Knutson, 2009) and the limbic “reward” system (Haber and Knutson, 2009). More specifically, human neuroimaging and primate neurophysiology have shown that goal-directed learning recruits large-scale cortical circuits including the lateral and medial prefrontal cortex (lPFC and mPFC), posterior parietal cortex, orbito-frontal cortex (OFC), temporal gyrus and visual cortex, and the temporal parietal junction (Tanaka, Balleine and O’Doherty, 2008; Walton *et al*., 2010; Liljeholm *et al*., 2011, 2013; Jocham *et al*., 2016; Averbeck and Costa, 2017; Bartolo and Averbeck, 2020; Norton and Liljeholm, 2020; Averbeck and O’Doherty, 2021; Morris *et al*., 2022).

Goal-directed learning is rooted in the balance between reward maximisation and information seeking processes (Cohen, McClure and Yu, 2007; Frank *et al*., 2009; Badre *et al*., 2012; Gottlieb *et al*., 2013; Mehlhorn *et al*., 2015; Schwartenbeck *et al*., 2019; Cockburn *et al*., 2022). Reward maximisation is formalised using associative (Rescorla, 1991) and reinforcement learning models (Watkins and Dayan, 1992; Sutton and Barto, 2018), where agents learn by maximising cumulative rewards. Update of action values is driven by reward prediction error (RPE) signals, which indicate whether an outcome is better or worse than expected. At the neural level, RPE signals are encoded by neurons in the midbrain, ventral striatum and ventromedial prefrontal cortex (vmPFC) (Frank, Seeberger and O’reilly, 2004; O’Doherty *et al*., 2004; Pessiglione *et al*., 2006; Schultz, 2006; D’Ardenne *et al*., 2008; Palminteri *et al*., 2015; Gueguen *et al*., 2021). In parallel, information seeking processes support learning by reducing the uncertainty in the causal relation between actions and outcomes, through signals quantifying the degree of surprise and information gain (Friston *et al*., 2015; Schwartenbeck *et al*., 2019). Information gain (IG) is formalised in model-based RL algorithms or Bayesian models (Yu and Dayan, 2005; Gläscher *et al*., 2010; Friston *et al*., 2015; Faraji, Preuschoff and Gerstner, 2018; Gottlieb and Oudeyer, 2018; Schwartenbeck *et al*., 2019; Liakoni *et al*., 2021; Modirshanechi *et al*., 2023) by Bayesian surprise (Itti and Baldi, 2009; Baldi and Itti, 2010), which quantifies how much the agent’s belief changes given new observation. Classical associative learning models formalise this process using the concepts of “surprisingness” or “associability” of events (Mackintosh, 1975; Pearce and Hall, 1980; Courville, Daw and Touretzky, 2006). At the neural level, IG has been mapped to the activity of distributed brain networks including the middle frontal gyrus, the insula and the intraparietal sulcus (Gläscher *et al*., 2010; Lee, Shimojo and O’Doherty, 2014; Fouragnan, Retzler and Philiastides, 2018; Liakoni *et al*., 2022; Modirshanechi *et al*., 2023). Neural correlates of subjective value of information in instrumental settings (i.e., information that can be used to guide future actions and future outcomes) have been observed in a distributed network including the ventral striatum, the vmPFC, the middle and superior frontal gyrus (i.e., the dorsolateral prefrontal cortex dlPFC), and posterior cingulate cortex (Kobayashi and Hsu, 2019; Kobayashi *et al*., 2021; Kobayashi and Kable, 2024).

Accumulating evidence suggests that the brain encodes update signals supporting reward maximisation and information gain during goal-directed learning. However, the role of cortico-cortical functional interactions underlying goal-directed learning signals remains elusive. To address this issue, we used information decomposition techniques to investigate whether cortico-cortical functional interactions, defined as statistical relationships between the activity of different cortical regions (Panzeri *et al*., 2022), encode learning signals for reward maximisation and information gain. In particular, we tested two hypotheses about the nature of functional interactions. First, we tested whether brain circuits encode information about cognitive processes by means of redundant and synergistic functional interactions (Varley, Sporns, *et al*., 2023; Luppi *et al*., 2024). Redundant encoding enhances robustness to perturbations by ensuring that critical information is consistently duplicated, and also facilitates the downstream readout of information (Valente *et al*., 2021; Combrisson *et al*., 2024). However, it may lead to inefficiencies by consuming excessive resources and limiting overall encoding capacity. Synergistic encoding enhances flexibility by supporting the emergence of novel information and richer representations by means of complementary encoding. Yet, it is less robust, because it relies on the precise integration of multiple inputs to generate new, emergent information. How the brain trades off redundant and synergistic encoding remains unaddressed.

The second hypothesis is that cognitive functions emerge from higher-order brain interactions, beyond pairwise relations (Martignon *et al*., 2000; Yu *et al*., 2011; Shahidi *et al*., 2019; Chelaru *et al*., 2021; Panzeri *et al*., 2022; Varley, Pope, Maria Grazia, *et al*., 2023a). Generally, higher-order functional interactions have the potential to link multiple neurons or brain regions in complex ways, and they can potentially generate emergent phenomena such as information integration, cognitive flexibility, and broadcast information (Schneidman *et al*., 2006; Köster *et al*., 2014; Chelaru *et al*., 2021; Panzeri *et al*., 2022; Santoro *et al*., 2024). The two hypotheses (tradeoffs of synergy/redundancy and high-order interactions) are not mutually exclusive and may allow an efficient encoding of information revelation for learning. More specifically, we expected to identify a distributed synergistic encoding and broadcasting of learning signals over multiple brain areas of the goal-directed and reward circuits in the prefrontal cortex.

Here, we used an experimental task design that manipulates learning and induces highly reproducible explorative strategies across participants and sessions. We combined information decomposition techniques based on partial information decomposition (Williams and Beer, 2010; Wibral *et al*., 2017; Lizier *et al*., 2018) and source-level high-gamma activity (HGA, from 60 to 120Hz) from magnetoencephalography (MEG) to test whether cortico-cortical interactions over learning circuits encode and broadcast reward prediction-error and information gain. We found that information gain is encoded in a distributed network including the visual, parietal, lateral prefrontal and ventro-medial and orbital prefrontal cortices. Interestingly, information gain was encoded in both redundant and synergistic cortico-cortical functional interactions. These functional interactions displayed complementary local-*versus-*distributed properties, synergistic interactions begin more prone for long-range interactions. Synergistic interactions displayed higher-order behaviours with a key role played by the prefrontal reward circuit, which acted as a receiver in the network. Overall, we suggest that higher-order synergistic interaction play a role in encoding and propagating learning signals in the brain, with a pivotal role played by the limbic cortical regions.

## RESULTS

### Behavioural scores, exploration strategy and learning signals

The experimental setup was structured to control the exploratory phase and guarantee consistent performance across sessions and individuals (Brovelli et al., 2008). The correct stimulus-response associations were not set *a priori*. Instead, they were assigned as learning proceeds in the task (Fig. 1b). Participants were asked to discover the associations existing among stimuli (coloured circles) and actions (finger movement). The task was controlled in such a way that the first presentation of each stimulus was always followed by an incorrect outcome, irrespective of subject’s choice (Fig. 1b, trials 1 to 3). On the second presentation of stimulus S1, any new untried finger movement was considered as a correct response (trial 4 in Fig. 1b). For the second stimulus S2, the response was defined as correct only when the subject had performed 3 incorrect finger movements (trial 9 in Fig. 1b). For stimulus S3, the subject had to try 4 different finger movements before the correct response was found (trial 14 in Fig. 1b). In other words, the correct response was the second finger movement (different from the first tried response) for stimulus S1, the fourth finger movement for stimulus S2, the fifth for stimulus S3. This task design assured a minimum of eight incorrect trials during acquisition (1 for S1, 3 for S2 and 4 for S4).

**Figure 1.**
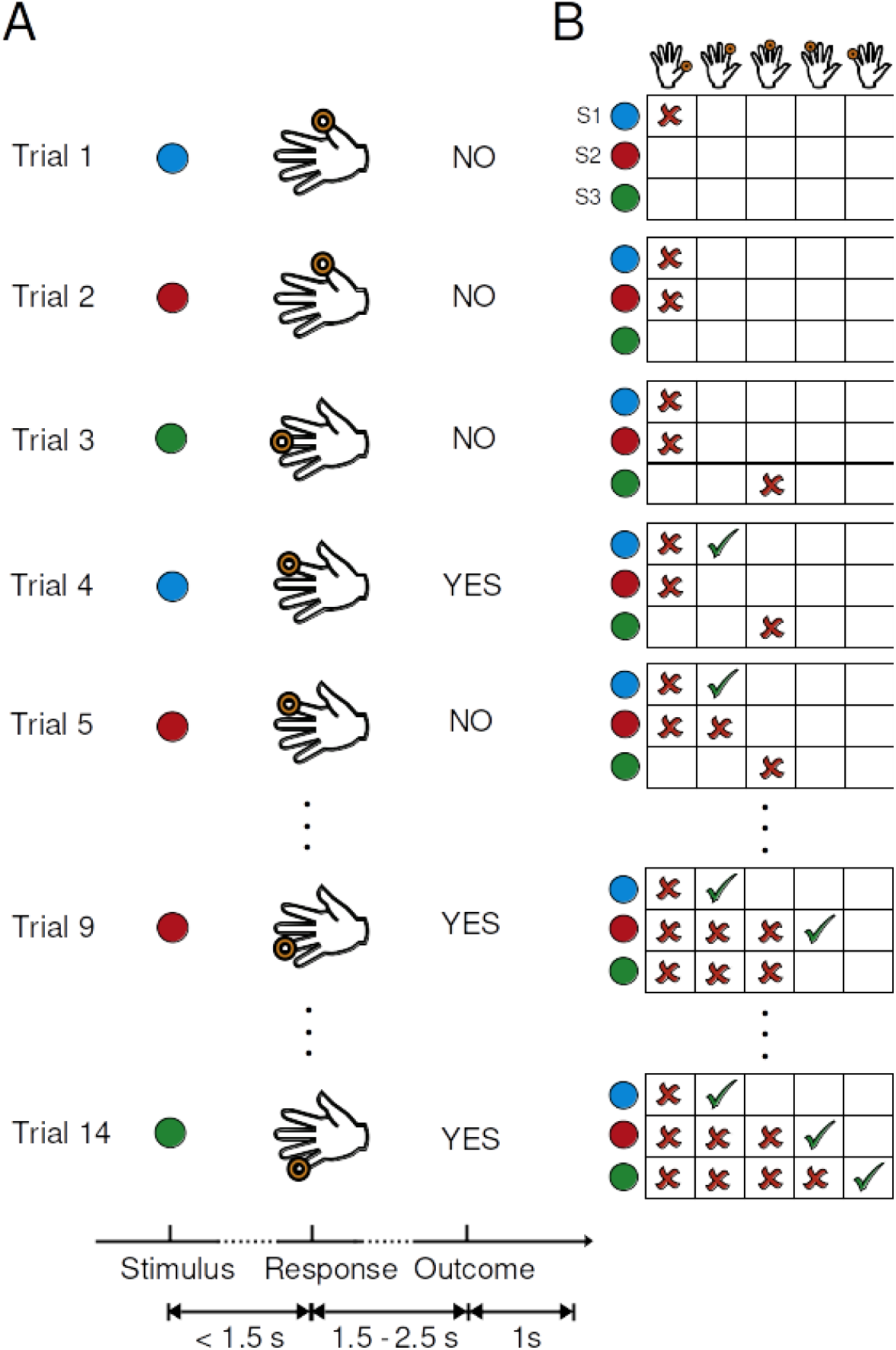
Experimental design and schematic representation of an ideal participant performing the learning task. **(A)** On each trial, subjects were presented a coloured circle to which they had to respond within 1.5 s. The outcome image was presented after a variable delay ranging from 1.5 to 2.5 s (randomly drawn from a log-normal distribution). The outcome image informed the subject whether the response was correct, incorrect, or too late (if the reaction time exceeded 1.5 s). **(B)** Matrix of all the possible stimulus–response combinations, updated according to the exemplar learning session in Fig. 1A for an ideal participant. A red cross and a green tick-mark refer to incorrect and correct stimulus–response sequences, respectively. The first presentation of each stimulus was always followed by an incorrect outcome, irrespective of the motor response (from trials 1 to 3). On the second presentation of the stimulus S1 (the blue circle), any untried finger movement was always followed by a correct outcome (trial 4). The correct response for S2 and S3 (red and green circles, respectively) was found after 3 and 4 incorrect finger movements (at trials 9 and 14, respectively).

Each participant performed four sessions of 60 trials each (total of 240 trials). Each session included three colours randomly presented in blocks of three trials. The average reaction time was 0.504s ± 0.04s (mean ± s.e.m.). The total number of error trials prior to the first correct outcome was 9.775 ± 0.38 (mean ± s.e.m.), thus indicating a low variability across participants and sessions. This indicated that the task produces reproducible behavioural performances across sessions and subjects, as observed in a previous study (Brovelli *et al*., 2008). The total number of errors after the first correct outcome was 4.15 ± 0.34, and they represented approximately 9% of the trials during the exploitative phase.

For what concerns the behavioural strategies, participants employed a “tree-search” heuristic, characterised by a tendency to repeat the same finger movement across trials until the correct response was found. This strategy is referred to as “lose-stay”, where agents maintain the response when not rewarded. Given that stimuli were pseudo-randomised in blocks of three trials, we computed the probability of repeating a given choice within a block of trials. This probability quantified the likelihood of a directed exploration, as opposed to random. For the first block (trials 1 to 3), the adoption of a lose-stay strategy occurred in 67.5% ± 6%, whereas in the second block of three trials (trials 4 to 6), it occurred in 37.5% ± 4.5%.

If the participants had employed a random exploration strategy, the proportion of consecutive trials with identical actions would be equivalent to the ratio of the total number of three-trial patterns with matching actions to the number of possible ordered triplets derived from all combinations of 3 distinct actions chosen from 5 available options (m = 60 combinations). Therefore, the likelihood of encountering a sequence of trials with the same action due to chance alone would be 8.3%. Our results demonstrate that participants used a directed exploration strategy during learning and rapidly attained learning and proficiency during each learning session.

A Q-learning model (Watkins & Dayan, 1992; Sutton & Barto, 1998) was fitted to the behavioural data to estimate changes in stimulus-action-outcome probabilities and learning signals over trials. The fitted model provided accurate predictions of learning curves, defined as the the probability of correct responses over trials (Suppl. Figure 1). Reward prediction errors (RPE) and information gain (IG) signals were extracted from the model. While RPE represents the discrepancy between the received and the anticipated outcome, IG measures the distance between the probability distributions of actions after and before the outcome. RPE embodies a scalar learning signal, fluctuating positive or negative based on whether reality surpasses or falls short of expectations, respectively. IG captures the magnitude of information conveyed by an outcome, and the amount of update in the associative model. We should note that RPE and IG do not covary monotonically, but display a U-shape relationship. Errors (i.e., negative RPE) and successes (i.e., positive RPE) are both associated with positive IG. As learning advances, both RPE and IG tend to zero (Suppl. Fig. 2).

### Brain areas encoding information gain signals during learning

To characterise both the spatial organisation and temporal dynamics of brain areas encoding learning signals, we performed time-resolved mutual information analyses between the across-trials high-gamma activity (HGA) and outcome-related learning signals, such as reward prediction errors (RPE) and information gain (IG). Group-level random-effect analyses, combining permutation tests and cluster-based statistics (see Methods), were used to identify temporal clusters displaying significant encoding of learning signals and HGA. We did not find any cortical area significantly encoding (*p*_*cluster*_ < 0. 05) RPEs. On the other hand, a large-scale network comprising visual, temporal, parietal and frontal areas displayed significant encoding of IG (Fig. 2). Forty brain regions were found to encode significantly IG, approximately 49% of the brain regions in the atlas. The strongest and earliest response was observed in the bilateral occipital regions, with a peak centred on the primary visual areas (VCcm and CMrm) of *MarsAtlas* (see top middle inset of Fig. 2), corresponding to Brodmann areas (BA) 17, 18, 19 and 39. Significant encoding was also observed in the left medial and superior parietal cortices (SPCm and PCm), corresponding to BA 7 and 31, respectively. In the temporal lobe, the right temporal cortices displayed significant encoding of IG. In the right frontal lobe, the right premotor areas (PMdl and PMdm), BA 6 and 8, and the right dorsolateral prefrontal cortices (PFcdm, PFcdl, Pfrdli and Pfrdls, PFdls, PFrdl and PFrdll), roughly corresponding to BA 45, 46, 27 and the most posterior parts of BA 9 and 10. Finally, the bilateral activation of the ventromedial prefrontal cortex (PFCvm, BA 32/10/11), orbitofrontal areas (OFCv, OFCvl and OFCvl, roughly BA 10/11/47) and the insula were found to display significant encoding of IG. Supplementary Figure 3 shows the same results shown in Fig. 2, but depicted on an inflated brain plot using *MarsAtlas* parcellation (Auzias et al., 2017).

**Figure 2.**
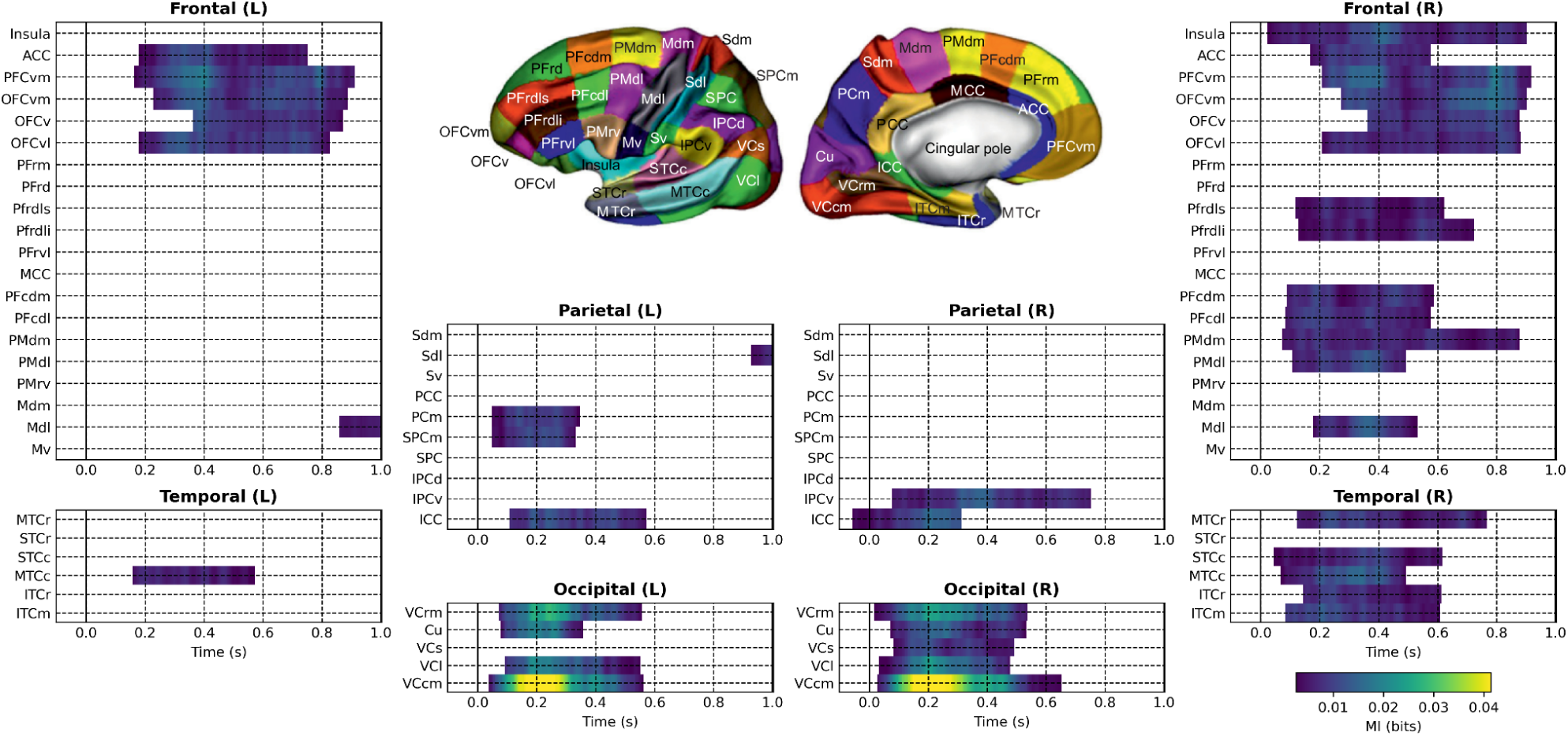
**Group-level statistics and** local spatio-temporal correlates of information gain. Regions-of-interest (ROI) of the MarsAtlas (Auzias et al., 2017) are shown in the inset in the top-central part of the figure. ROIs are grouped in four panels according to the different lobes (occipital, parietal, temporal and frontal) and hemisphere (left and right panels). In each panel, colored areas indicate the temporal clusters for which there is a significant encoding of information gain. The p-values for each cluster accounted for multiple comparisons and they were threshold to 0.05. Colormap shows the value of the mutual information (in bits) between the HGA and IG. Time in seconds is on the x-axis with respect to outcome onset (time 0). A large-scale network encodes IG and it recruits the occipital, parietal, right temporal, right dorsolateral prefrontal areas, ventromedial prefrontal cortex and, orbitofrontal areas.

Overall, IG was encoded in a set of eight cortical subnetworks (or clusters) including bilateral visual areas (the “L Vis” cluster included the left VCcm, VCl, Cu and VCrm; cluster “R Vis” included the right VCcm, VCl, Cu, VCrm and VCs), the right temporal regions (cluster “R Temp” included the right ITCm, ITCr, MTCc, STCc and MTCr), the left superior parietal area (cluster “L Par” included the left PCm and SPCm), the right motor and premotor areas (cluster “R MotPM” included the right Mdl, PMdl and PMdm), the right dorsolateral prefrontal cortices (cluster “R dlPFC” included the right Pfrdli, Pfrdls, PFcdm and PFcdl) and the bilateral ventromedial prefrontal and orbitofrontal areas (cluster “R vmPFCOFC” included the right OFCvl, OFCv, OFCvm, PFCvm, whereas the cluster “L vmPFCOFC” included the left OFCvl, OFCv, OFCvm and PFCvm). Such grouping was based on anatomical constraints and belonging to the same lobe or anatomical region.

In order to characterise the time course of learning-related signals encoding IG, we averaged the mutual information across areas within each height cluster of cortical regions displaying significant effects (Fig. 3). The aim was to extract an average time course for each anatomical cluster. The average time courses are shown in Fig. 3. The visual (Fig. 3A-B), temporal (Fig. 3C) and parietal (Fig. 3D) clusters displayed a time course characterised by a fast and transient increase in MI and peaking around 0.2-0.3s after outcome onset.

**Figure 3.**
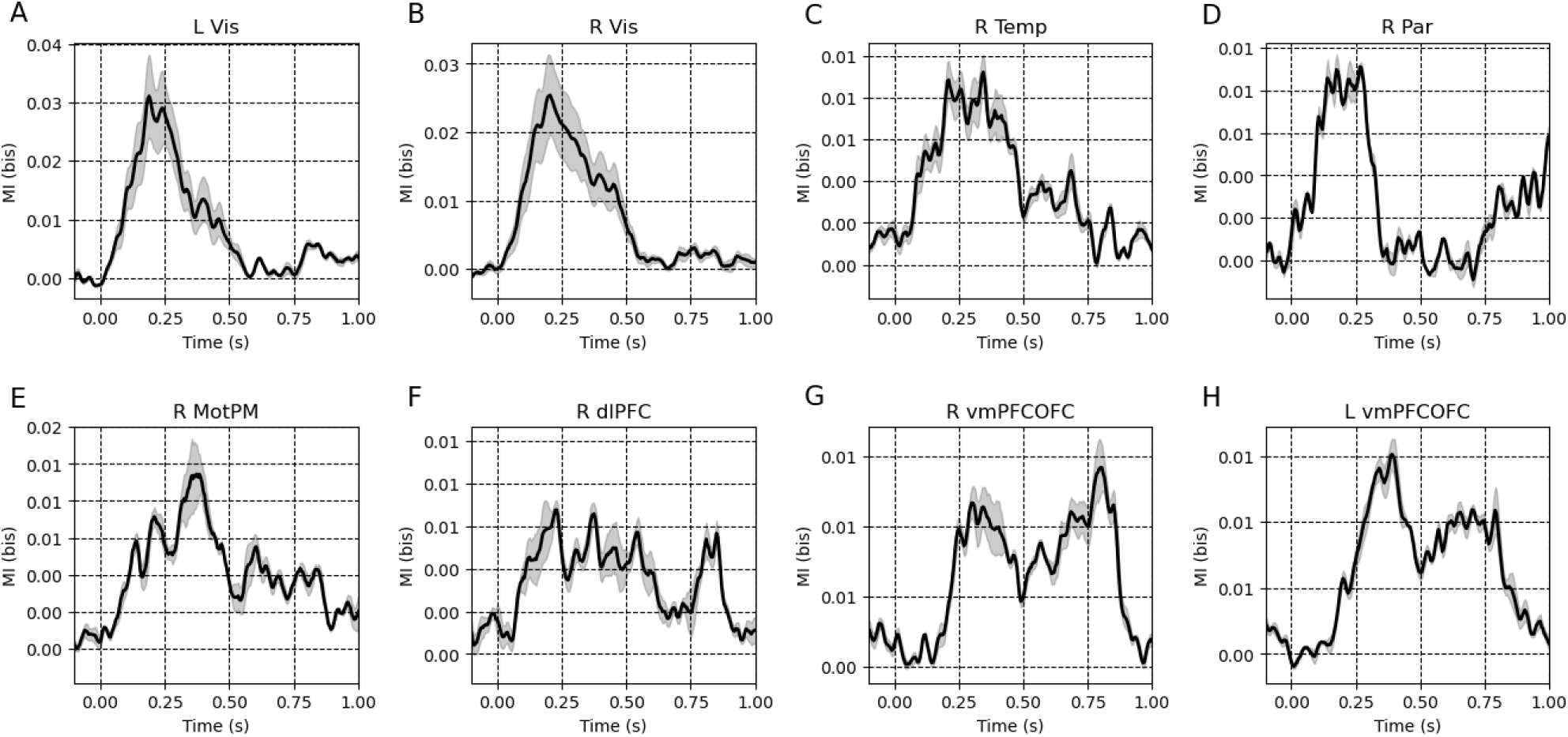
Time courses of mutual information values between HGA and IG for the eight clusters of cortical areas. The time axis indicates the time from the outcome presentation in seconds. A, B) Left and right visual areas, respectively; C) Right temporal area; D) Right parietal area; E) Right motor and premotor areas; F) Right dorso-lateral prefrontal cortex; G, H) Right and left ventromedial prefrontal and orbitofrontal cortices respectively. Time in seconds is on the x-axis with respect to outcome onset. Shaded areas are standard errors of the mean computed across ROIs within each cluster.

The encoding in temporal regions peaked around 0.3s after outcome onset (Fig. 3C) and returned to based around 0.8s after outcome onset. The premotor and dorsolateral regions of the right hemisphere peaked approximately from 0.3 to 0.4s after outcome and they were characterised by a slower return to baseline (Fig. 3E). Finally, the vmPFC and OFC regions were characterised by an onset at approximately 0.2s and an elevated encoding lasting until almost 1s (Fig. 3F to 3H).

To differentiate complementary spatio-temporal patterns of encoding of IG, we performed non-negative matrix factorisation (NMF) of the MI values across clusters. The rationale was to decompose the MI time courses into interpretable temporal components and corresponding spatial weights. Figure 4A shows the variance explained by the first ten components of the non-negative matrix factorisation. Based on the so-called the elbow method, which is a standard heuristic for determining the number of components in a dataset, we studied the first four components whose loadings are depicted in Fig. 4B. . Figure 4C shows the variance explained of the top four components, normalized by their total variance (in colour) for each brain area (rows). The first component (in blue) modelled the fastest and most transient encoding of IG and it is mainly observed in the visual and parietal areas (blue bars in Fig. 4C). The orange and green components occurred later in the trial, peaking around 0.35 and 0.45s after outcome onset, recruiting primarily the temporal, premotor and dorsolateral areas. Finally, the fourth component (in red) captured the slowest component that emerged around 0.5s and lasted approximately 0.4s. This latest component mainly recruited the ventromedial PFC and OFC cluster. Overall, these results show that the encoding of IG recruits a large-scale network of cortical clusters with overlapping temporal and spatial dynamics.

**Figure 4.**
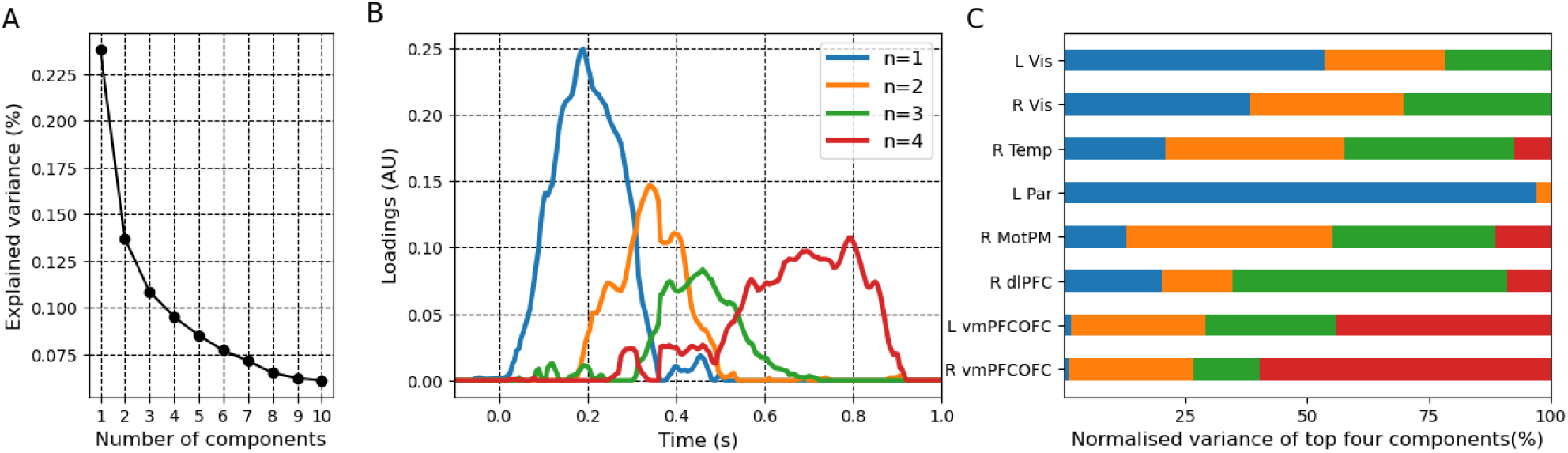
(A) Variance explained by each component of the non-negative matrix factorisation. (B) First four loading of the non-negative matrix factorisation. The color of the loading indicates the component number (see legend). The time axis indicates the time from the outcome presentation in seconds. (C) Variance explained of the top four components, normalized by their total variance. Each row shows the percentage of variance explained by each component across all brain regions. The color of each bar indicates the component number (see legend in panel B).

### Cortico-cortical interactions mediating information gain

We then tested the hypothesis that information gain is encoded at the level of cortico-cortical interactions, in addition to local activations. To do so, we exploited recent advances in information theory that allow isolating the contribution of network-level interactions from those carried at the node level. We thus decomposed the information carried by pairs of brain regions about IG into redundant (i.e., shared or common between regions) and synergistic (i.e., requiring the joint combination) terms, as formalised by the Partial Information Decomposition (PID) framework (Williams and Beer, 2010; Lizier et al., 2018). The Partial Information Decomposition (PID) framework enables the breakdown of multivariate mutual information between a set of predictor variables and a “target” variable into distinct, non-negative components. These components quantify the unique, redundant, and synergistic contributions of the predictors in conveying information about the target variable. In our analysis, we measured the redundant and synergistic information provided by pairs of brain regions (predictors) regarding IG (the target variable). The predictor variables consisted of the high-gamma activity (HGA) recorded across trials from two brain regions, while the target variable represented the trial-by-trial evolution of IG. Redundant information captures the shared contribution of both brain regions to IG, meaning the information that regions share. In contrast, synergistic information reflects the additional information carried only when both regions are observed together, which cannot be obtained from either one individually.

Significant redundant information about IG was observed across multiple brain regions (Fig. 5A). Strongest redundant connectivity was observed earlier over bilateral visual areas during the first 0.2s after outcome presentation (left panel of Fig. 5A). Redundancy connectivity then spread over the right temporal lobe, right motor and premotor cortices, and vmPFC and OFC in the time interval between approximately 0.2 and 0.8s. Redundant connectivity over the vmPFC - OFC cluster were long lasting and faded away approximately 0.8 to 1s after outcome (rightmost panel of Fig. 5A). Interestingly, we observed strong synergistic interactions between specific pairs of brain regions (Fig. 5B), indicating that these regions jointly contribute information about the target variable in a way that neither region provides independently. Synergistic connectivity appeared to link distant areas of the visual system, the right temporal areas, the right dlPFC and the left vmPFC and OFC. The peak of interaction strength occurred from 0.2 to 0.5s after the outcome presentation. Redundant connectivity appeared to predominantly recruit pairs of brain regions with the same region or cluster, whereas synergistic connectivity appeared to be stronger across systems. In order to verify such an effect, we computed the average redundant and synergistic connectivity between pairs of brain regions within and across anatomical clusters in the visual, temporal, motor and premotor, dlPFC and vmPFCOFC clusters. The visual cluster included the left and the right VCcm, VCl, Cu and VCrm areas; the temporal cluster included the right ITCm, ITCr, MTCc, STCc and MTCr; the motor and premotor cluster included the right Mdl, PMdl and PMdm; the dorsolateral PFC included the right Pfrdli, Pfrdls, PFcdm and PFcdl; and finally the vmPFCOFC region included the bilateral OFCvl, OFCv, OFCvm, PFCvm, OFCvl, OFCv, OFCvm and PFCvm. Indeed, we found that redundant connectivity showed stronger average value across pairs of brain regions within the same module (Fig. 6A), whereas synergistic coupling showed a stronger effect across modules (Fig. 6B).

**Figure 5.**
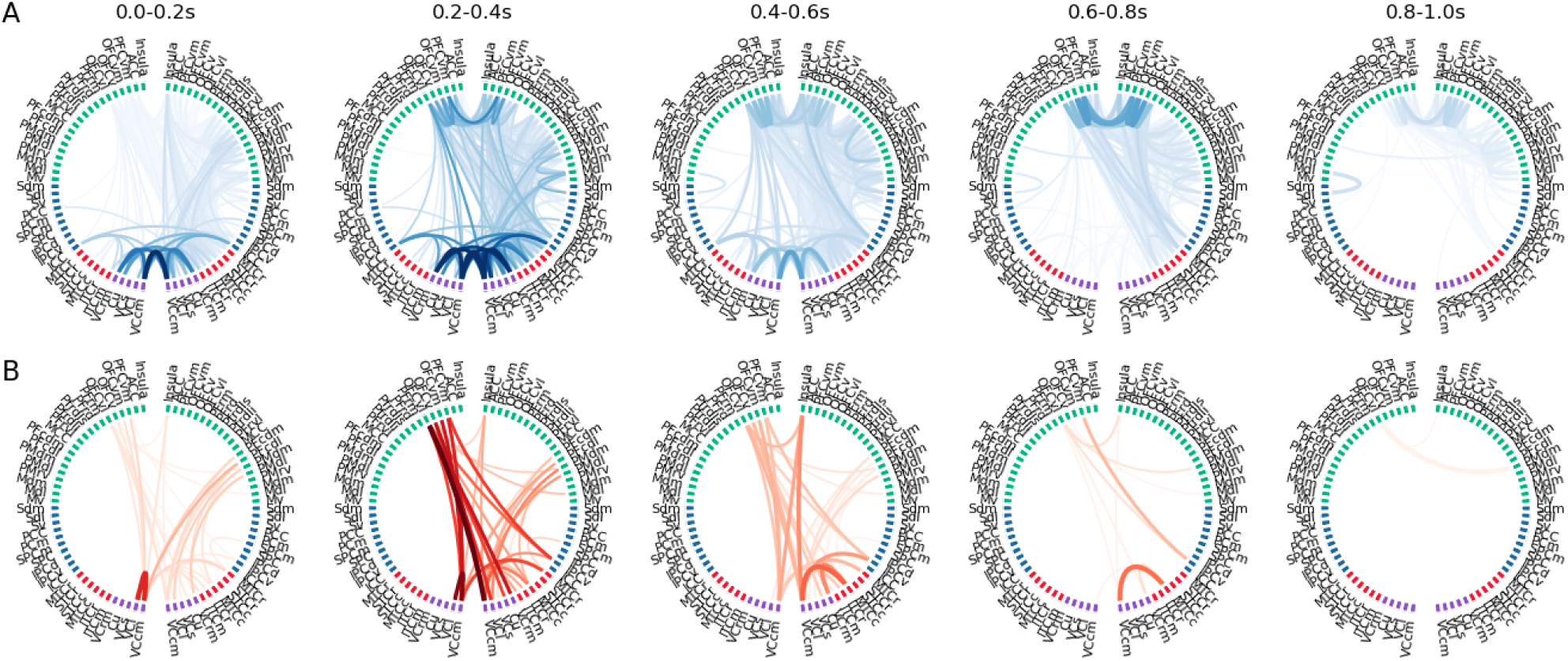
Redundant (A) and synergistic (B) pairwise interactions significantly carrying information about information gain signals. Each panel is the average value in 0.2s time windows aligned on outcome presentation. The labels of the regions of interest plotted in a circle are identical to those of Fig. 2. The position of each brain region over the cortical sheet and lobe can be seen in the inset of Fig. 2. The p-values for each cluster accounted for multiple comparisons and they were threshold to 0.05. The color of each node indicates its association with a particular lobe (visual in purple, temporal in red, parietal in blue, and frontal in green).

**Figure 6.**
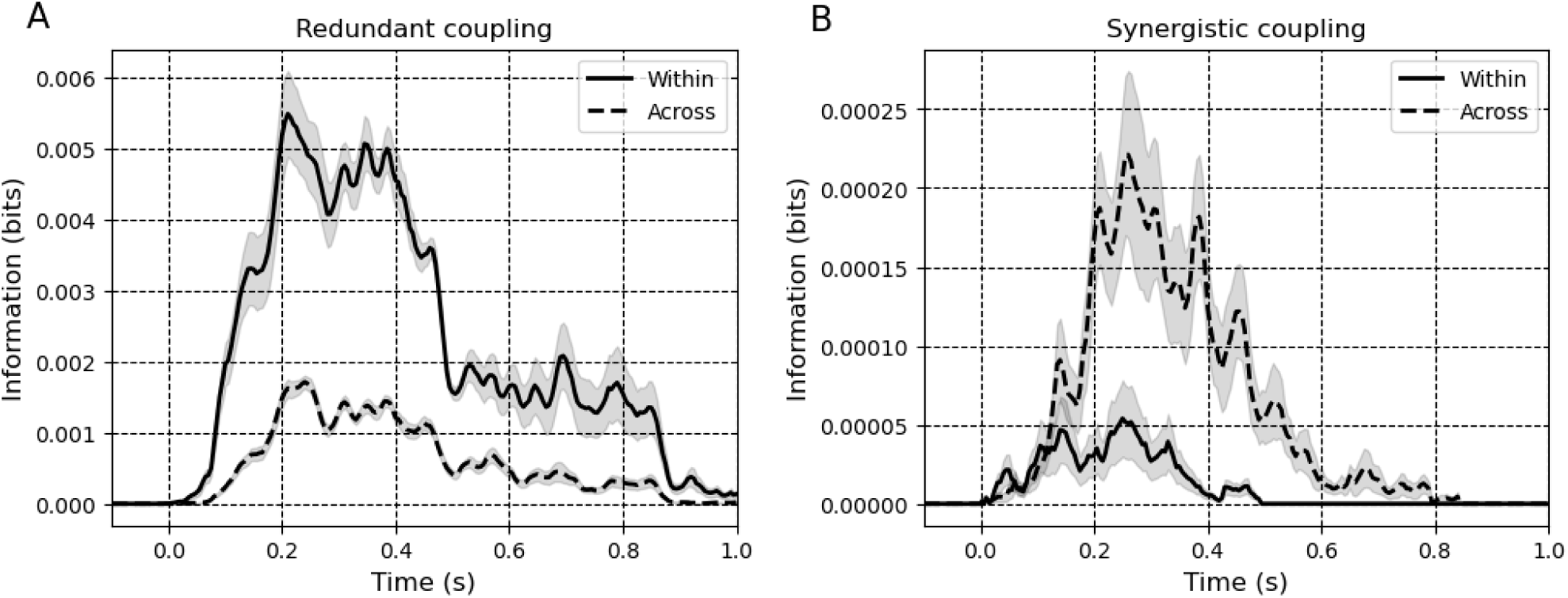
Average redundant (A) and synergistic (B) connectivity computed over pairs of brain regions being part of the same (within) or different (across) cluster. The “within-cluster” connectivity is represented by a solid line, while the “across-cluster” connectivity is depicted using a dotted line. Mean and standard error of the mean are plotted as solid line and shade area, respectively. The time axis indicates the time from the outcome presentation in seconds. Please note that the scales are different across panels (A) and (B).

By definition (Eq. 9), redundant functional interactions represent the smallest amount of information about learning-related variables that is already present in each individual brain area. In other words, they quantify the overlap in information shared by brain regions, reflecting the common contribution that each region independently provides about the learning process. On the other hand, synergistic connectivity reflects functional interactions that arise only through the combined activity of multiple brain areas, rather than being explained by the individual contributions of each region alone. Significant synergistic functional interactions therefore reveal the presence of collective behaviour encoding IG that emerge from the coordination of distant brain areas and circuits. This highlights the important role of collective and synergistic processes beyond the contribution of individual brain areas and redundant information encoding.

### Higher-order cortico-cortical interactions and information gain

Next, we investigated whether information relevant to cognitive functions, such as IG, is encoded in a distributed manner across multiple brain regions, extending beyond simple pairwise interactions to higher-order dynamics (Martignon et al., 2000; Yu et al., 2011; Shahidi et al., 2019; Chelaru et al., 2021; Panzeri et al., 2022; Varley, Pope, Maria Grazia, et al., 2023). Building on recent studies that emphasize higher-order correlations beyond pairwise interactions as a potential foundation for emergent phenomena (Gatica et al., 2021; Herzog et al., 2022; Varley, Pope, Maria Grazia, et al., 2023), we examined whether cortico-cortical interactions encode IG through higher-order co-modulations. We anticipated a central role for brain regions involved in reward and goal-directed circuits, such as the vmPFC, OFC, and dlPFC, potentially carrying higher-order synergistic information through interactions among multiple regions.

To do so, we studied higher-order relations between the eight cortical clusters encoding IG (Fig. 3 and 4). Since each cluster independently encodes information gain, they are also expected to contribute to redundant information at higher-order relational levels. However, higher-order synergistic connectivity—representing information that emerges from collective patterns beyond pairwise relationships—requires further investigation. Cluster-based statistical analyses and correction for multiple comparisons (across all pairs and multiples across all orders) were performed to identify significant higher-order encoding of IG. Statistical analyses revealed a total of 15 pairs, 7 triplets and 4 quadruplets to be encoding significantly IG (Fig. 7A). Higher-order synergistic co-modulations carrying information about IG were found up to order 4. Given that we analysed 8 clusters, the percentage of significant pairwise links, triplets and quadruplets was 53%, 12.5% and 6%, respectively. We next asked whether higher-order functional interactions were associated with hypergraph or simplicial complex representations. As a reminder, a hypergraph is a description of higher-order interactions, consisting of k nodes representing a k-way interaction between the nodes. A simplicial complex is a special case of a hypergraph, where all possible (k-1)-way interactions among that same set of nodes occur (Battiston *et al*., 2020, 2021). In other words, hypergraphs without lower-level interactions may reflect a stronger “marker” of emergent collective behaviour, because it would lack lower level components. We found that 6 out of 7 triplets were composed of significant pairwise links, and therefore reflected simplicial complexes (Fig. 7A). Only one triplet was composed of nodes that did not display all pairwise interactions, therefore reflecting a stronger emergent form of collective behaviour (L Vis - R MotPM - L vmPFCOFC). The four quadruplets were not composed of all lower order triplets. This can be observed in Suppl. Figure 4, which depicts the same results but using hypergraph representations. Overall, these results indicate that IG is encoded in a distributed fashion through synergistic interactions that extend beyond individual brain regions, revealing genuine higher-order synergy.

**Figure 7.**
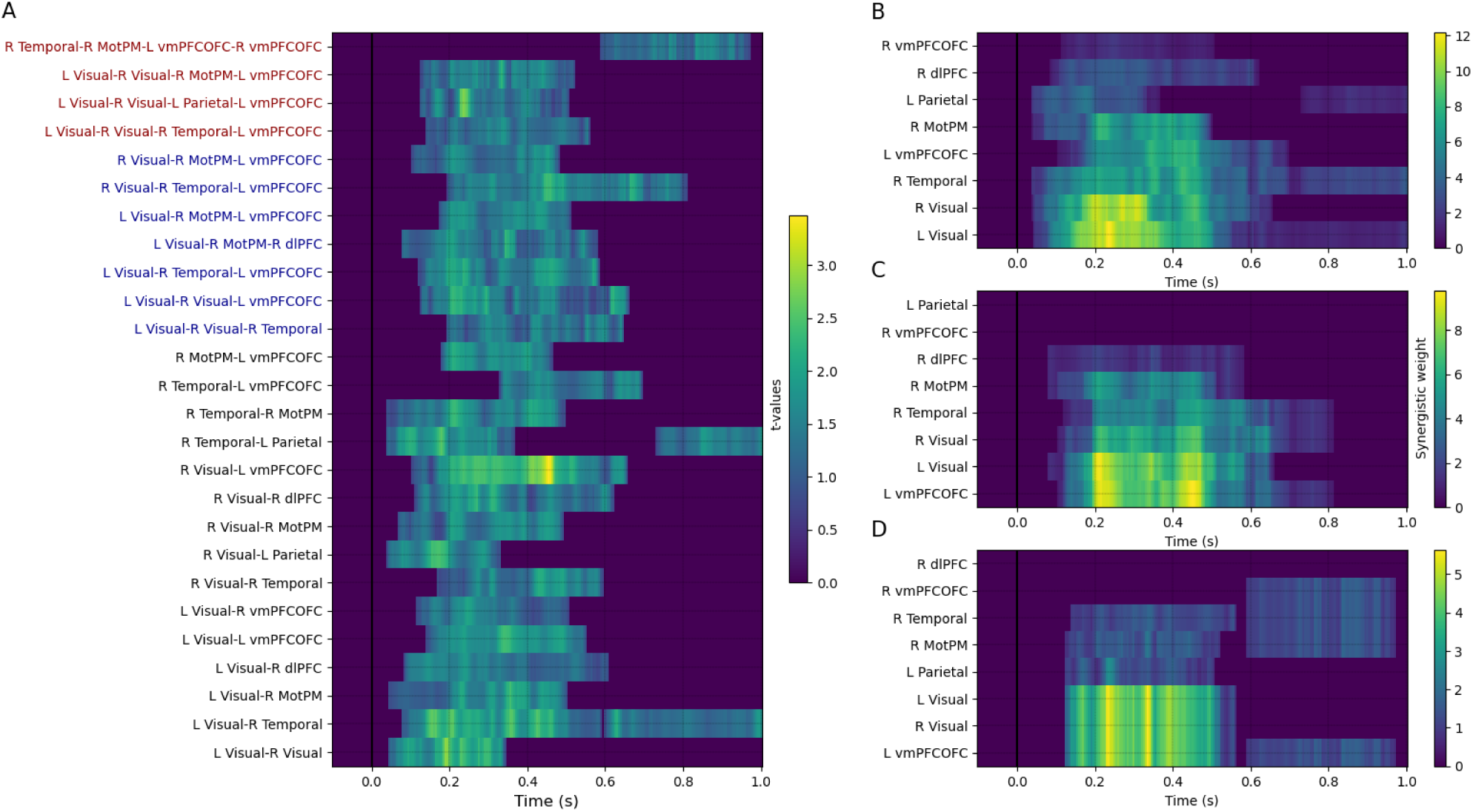
Synergistic higher-order interactions between cortical clusters encoding information gain (IG). (A) Time course of t-values showing significant encoding of IG at the level of pairs of clusters (black labels), triplets (blue label) and quadruplets (red labels). We then computed the synergistic weight, defined as the sum of the synergistic information for each cluster. This value reflects the amount of contribution in synergistic encoding across scale. (B) Synergistic weight for clusters and for pairwise interactions. (C) and (D) show cluster weight for triplets and quadruplets, respectively. The time axis indicates the time from the outcome presentation in seconds. In panels (B), (C), and (D), clusters are ranked in ascending order based on their maximum synergistic weight value.

Significant temporal clusters were mainly observed over the interval between 0.1s and 0.6s, and no clear differentiation between pairwise and 3rd- or 4th-order interactions. In order to better characterise the participation of clusters in each multiple, we computed the synergistic weight defined as the summed synergistic connectivity. Figure 7B shows the nodal synergistic weight at the pairwise level, highlighting that the bilateral visual clusters exhibit the highest values. However, at higher orders, the visual clusters were accompanied by the left vmPFC-OFC, which dominated both triplet (Fig. 7C) and quadruplet (Fig. 7D) interactions. This indicates that the left vmPFC-OFC, a key region in the reward circuits, gained prominence at higher orders of synergistic interactions.

### Cortico-cortical information transfer encoding learning signals

Previous results highlighted a time-lagged encoding of IG from visual to prefrontal areas, suggesting a feedforward broadcasting of IG signals to prefrontal regions (Fig. 3 and 4). In order to test this hypothesis, we exploited a recently-developed information-theoretic measure termed Feature-specific Information Transfer (FIT). Feature-specific Information Transfer (FIT) quantifies how much information about specific features flows between brain areas (Celotto et al., 2023). FIT merges the Wiener-Granger causality principle (Granger, 1980; Brovelli et al., 2004; Bressler and Seth, 2011) with content specificity based on the PID framework (Williams and Beer, 2010; Lizier et al., 2018). FIT isolates information about a specific task variable Y (information gain) encoded in the current activity of a receiving neural population, which was not encoded in its past activity, and which was instead encoded by the past activity of the sender neural population. We used the FIT measure to quantify the broadcasting of IG between the eight clusters of brain regions. Four directional relations significantly encoded IG and broadcasted information to the vmPFC and OFC (Fig. 8). The temporal information iss aligned from the point of view of the receiving brain area. The IG-specific information transfer peaked around 0.35 to 0.4s after outcome onset, thus roughly corresponding to the first peak in HGA observed locally. Two patterns of interactions converge towards the left and right vmPFC-OFC, respectively. These “colliding” patterns included two higher-order synergistic multiplets, namely a triplet including the right visual and temporal regions with the left vmPFC-OFC, and a quadruplet including right visual and temporal regions with the bilateral vmPFC-OFC (Fig. 7).

**Figure 8.**
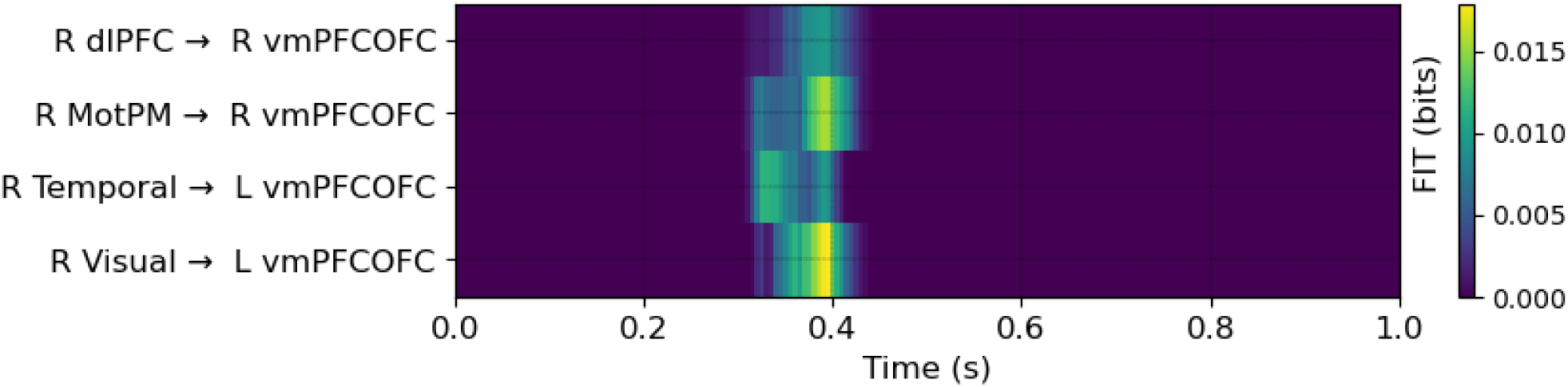
Feature-specific Information Transfer (FIT) encoding Information Gain (IG). Significant directional interactions broadcast information about IG primarily to the vmPFC-OFC. The time axis represents the time in seconds from the presentation of the outcome.

## Discussion

### Information seeking and directed exploration in goal-directed learning

Goal-directed learning is defined as the ability to learn the causal relationship between our behaviours and their outcomes; this supports the selection of actions according to expected outcomes, as well as current goals and motivational state (Tolman, 1948; Dickinson, 1994; Dolan and Dayan, 2013). Goal-directed learning relies on the balance between exploitation and exploration strategies (Cohen, McClure and Yu, 2007; Frank *et al*., 2009; Badre *et al*., 2012; Gottlieb *et al*., 2013; Schwartenbeck *et al*., 2019; Cockburn *et al*., 2022). A trade-off between choosing the most valuable option and sampling informative options is central for successful goal-directed learning and it is one of the main focus of reinforcement learning and Bayesian learning theories (Friston *et al*., 2015, 2017; Sutton and Barto, 2018). Exploration in humans has been shown to be based on a mixture of random and directed strategies (Wilson *et al*., 2014; Gershman, 2018). Random exploration is a deviation from the most rewarding option, and it is normally formalised using the softmax choice rule, which models the degree of randomness (decision noise). Directed exploration selectively samples options that are informative (information seeking), that are associated with the highest uncertainty.

Here, we have addressed the neural computations supporting information seeking during goal-directed learning in terms of information gain and directed exploration. In order to investigate directed exploration and information gain, we used a learning task previously developed for fMRI (Brovelli et al., 2008) that manipulates learning and induces reproducible explorative strategies across subjects and sessions. Learning was characterised by a directed exploration strategy based on a “tree-search” or “lose-stay” pattern, in which participants had a tendency to repeat the same action after incorrect outcomes and span all options in an ordered manner. Given that the correct action-outcome association was deterministic, such directed exploration strategy pertained only the initial phase of the task, up to the first correct outcome. Subsequently, participants performed a relatively low number of maintenant errors and engaged in an exploitative strategy. From the computational point of view, information seeking and exploration is formalised using model-based RL algorithms or Bayesian models (Yu and Dayan, 2005; Gläscher *et al*., 2010; Friston *et al*., 2015, 2017; Faraji, Preuschoff and Gerstner, 2018; Liakoni *et al*., 2021; Modirshanechi, Brea and Gerstner, 2022) and it has been proposed to play a general role in curiosity-driven learning (Gottlieb and Oudeyer, 2018; Schwartenbeck *et al*., 2019). The value of actions is not solely determined by the expected utility (or extrinsic value), but also by the informative (intrinsic, epistemic, or non-instrumental) value that is expected to be gained, which helps reduce uncertainty about environmental contingencies (Cohen, McClure, and Yu, 2007; Bromberg-Martin and Hikosaka, 2009; Tishby and Polani, 2011; Friston et al., 2015; White et al., 2019). Regarding the update of action values during learning, we propose that directed exploration involves at least two processes: i) updating the expected reward (extrinsic) value of an action via reward prediction errors (RPE), and ii) updating the (intrinsic or informative) value of an action using an information gain (IG) signal. IG signals track the trial-by-trial evolution of information gain during directed exploration. This information is used to reduce uncertainty about the causal relationship between actions and outcomes, thereby decreasing the agent’s uncertainty about the world. Within the taxonomy of surprise definitions, our study focuses on the subcategory known as “belief-mismatch surprise” (Modirshanechi, Brea, and Gerstner, 2022), which we refer to as information gain. A limitation of our study is the inability to link the neural results to alternative measures of information gain within the belief-mismatch surprise category, such as the minimization of free energy (Schwartenbeck et al., 2013).

### Distributed spatio-temporal cortical activations encoding information gain

To study how IG is encoded in the brain, we performed information theoretical analyses of single-trial and cortical-level high-gamma activity (HGA) from MEG. A network comprising visual, temporal, parietal and frontal areas was found to encode IG during goal-directed learning (Fig. 2). At the circuit level, the results suggest that information gain is encoded in the dorsal and ventral circuits of the goal-directed system (Averbeck B, 2020; Averbeck and O’Doherty, 2021). The dorsal circuit, comprising the dorso-lateral prefrontal cortex (dlPFC) and the inferior parietal cortices, is thought to primarily encode information relevant for actions and control behaviour to achieve goals. The ventral circuit, comprising the ventro-medial prefrontal cortex (vmPFC) and orbito-frontal cortex (OFC), is suggested to primarily learn to assign and update subjective values to potential outcomes and motivational value of objects in the environment, owing to visual input from the anterior temporal cortex and connections with the hypothalamus (Averbeck and Costa, 2017). Our results show that IG is encoded over both the ventral and dorsal circuits without a clear dissociation between circuits. Our findings confirm that IG and Bayesian surprise are represented in distributed brain networks including the middle frontal gyrus, the insula and the intraparietal sulcus (Gläscher *et al*., 2010; Lee, Shimojo and O’Doherty, 2014; Fouragnan, Retzler and Philiastides, 2018; Liakoni *et al*., 2022; Modirshanechi *et al*., 2023). Indeed, fMRI activity encoding belief updates (Bayesian surprise) has been observed in the midbrain and ventral striatum, in addition to the presupplementary motor area (pre-SMA), dorsal anterior cingulate cortex, posterior parietal cortex and lateral prefrontal cortex (O’Reilly *et al*., 2013; Nour *et al*., 2018). Our results complement fMRI studies showing correlates of information value in instrumental settings (i.e., information that can be used to guide future actions and future outcomes) in the ventral striatum, vmPFC, the middle and superior frontal gyrus (i.e., the dorsolateral prefrontal cortex dlPFC), and posterior cingulate cortex (Kobayashi and Hsu, 2019; Kobayashi *et al*., 2021; Kobayashi and Kable, 2024). Further work is needed to fully appreciate the relation between the processing of information about outcomes and action selection during information seeking.

At the hemispheric level, a right-lateralized activation was observed (Fig. 2 and Suppl. Fig. 3). This result may relate to established literature showing a right-hemisphere dominance for stimulus-driven shifts of spatial attention and target detection (Shulman *et al*., 2010; Corbetta and Shulman, 2011) and activation patterns observed in the ventral attentional network (Vossel, Geng and Fink, 2014). This result also resonates with associative learning models formalising attentional processes during learning using concepts such as “surprisingness” or “associability” of events (Mackintosh 1975; Pearce and Hall 1980; Courville et al. 2006). We propose that attentional processes, along with the neural circuits involved, contribute to information-seeking behavior, although the exact relationship between them remains unclear.

At the regional level, a key finding was the encoding of IG in vmPFC and OFC. This suggests a link to the reward circuits in limbic regions classically associated with the encoding of reward prediction error (RPE) signals (Frank, Seeberger and O’reilly, 2004; O’Doherty *et al*., 2004; Pessiglione *et al*., 2006; Schultz, 2006; D’Ardenne *et al*., 2008; Palminteri *et al*., 2015; Gueguen *et al*., 2021). Our results are in line with the notion of a “common currency” for information and reward values in the limbic system (Levy and Glimcher, 2012; Kobayashi and Kable, 2024). These areas are main targets of dopaminergic projections, which encode, in non-instrumental settings, expected advance information about future outcomes encoding errors in information prediction (information prediction error, reflecting the difference between obtained and expected information gain) (Bromberg-Martin and Hikosaka, 2009, 2011; Bromberg-Martin *et al*., 2024), in the OFC in human fMRI (Charpentier, Bromberg-Martin and Sharot, 2018) and in frontal EEG electrodes (Brydevall *et al*., 2018). The anterior cingulate cortex (ACC) and two subregions of the basal ganglia (the internal-capsule-bordering portion of the dorsal striatum and the anterior pallidum) signalling reward uncertainty and information-anticipatory activity (White *et al*., 2019) may additionally provide a substrate for common encoding of reward and information value. The underlying neural mechanisms are currently unknown, However, converging evidence supports that subpopulations of midbrain dopamine neurons encoding motivational values (Bromberg-Martin, Matsumoto and Hikosaka, 2010) may contribute to the observed activations in limbic regions. This may explain the assignment of value to uncertainty reduction, integrating the value of information with the value of physical rewards, and may provide a link with the neural circuits for information seeking and curiosity-driven behaviours (Monosov, 2024).

We suggest that the current results provide a comprehensive understanding of the role of ventromedial PFC and OFC in decision-making and information seeking. It is widely accepted that vmPFC/OFC plays a key role in learning and decision making by encoding outcome information and value (Padoa-Schioppa and Assad, 2006; Ballesta *et al*., 2020; Gardner *et al*., 2020; Gore *et al*., 2023). Moreover, during goal-directed exploration, fMRI activity in the rostral frontopolar cortex (RFPC) activity (Badre and D’Esposito, 2009) and frontal theta component in EEG have indicated a direct role in directed exploration (Cavanagh *et al*., 2012; Cavanagh and Frank, 2014), rather than random exploration (Daw, Niv and Dayan, 2005). Stimulation and inhibition of RFPC with direct current (tDCS) has shown to increase and decrease the frequency of exploratory choices (Raja Beharelle *et al*., 2015; Zajkowski, Kossut and Wilson, 2017). These studies put forward the idea that the RFPC is crucial for integrating information about past, present, and future payoffs to arbitrate between exploration and exploitation in human decision making. Our results suggest that a key computation supporting directed exploration and information seeking is the processing of information gain. Overall, the results provide strong support for the idea that information gain signals are encoded in the human brain. This mediates the brain’s capacity to integrate uncertainty-reducing information, as predicted by theories of information-seeking behavior (Yu and Dayan, 2005; Gläscher et al., 2010; Friston et al., 2015, 2017; Faraji, Preuschoff and Gerstner, 2018; Liakoni et al., 2021; Modirshanechi, Brea and Gerstner, 2022) and curiosity (Gottlieb and Oudeyer, 2018; Schwartenbeck et al., 2019). This also aligns with computational models that aim to combine reward maximization with information gain mechanisms (Friston et al., 2015; Schwartenbeck et al., 2019).

Our results provide insights into the temporal dynamics of information gain during directed exploration, an aspect that was lacking in previous neuroimaging and neurophysiological studies in primates. The strongest and earliest encoding of information gain occurred in occipital regions, followed by the parietal and temporal cortices, and the right frontal lobe including premotor and dorsolateral prefrontal cortices, and final activation of the ventromedial prefrontal cortex orbitofrontal areas and insula (Fig. 3). We statistically dissociated four complementary spatio-temporal patterns of HGA encoding IG (Fig. 4). The first component modelled the fastest and most transient encoding of IG recruiting the visual and parietal areas. The second and third component occurred later in the trial peaking around 0.35 and 0.45s after outcome onset and they recruited primarily the temporal, premotor and dorsolateral areas. Finally, the fourth component captured the slowest component that emerged around 0.5s and later approximated 0.4s, involving the ventromedial PFC and OFC cluster. Overall, our results demonstrated that the encoding of IG recruits a large-scale network of cortical clusters with overlapping temporal and spatial dynamics, spanning visual to limbic frontal areas.

### Redundant and synergistic functional interactions encode information gain

We next investigated whether and how information relevant for learning such as IG is encoded in cortico-cortical interactions. We observed that IG is encoded at the level of cortico-cortical interactions (Fig. 5), in addition to local activations (Fig. 2). We performed information decomposition analyses to quantify the information carried by pairs of brain regions, by dissociating the encoding of IG into redundant (i.e., shared or common between regions) and synergistic (i.e., requiring the joint combination) interactions. We observed that IG was encoded both by redundant and synergistic cortico-cortical co-modulations (Fig. 5). Strongest redundant connectivity was observed earlier over bilateral visual areas during the first 0.2s after outcome presentation and then spread over the right temporal lobe, right motor and premotor cortices, and vmPFC and OFC cluster. By definition (Eq. 9), redundant functional interactions equate the minimum of information carried locally. Redundant functional connectivity therefore reflects the shared information carried by co-modulations in HGA across areas (Fig. 5A). On the other hand, synergistic connectivity analysis reveals functional interactions that cannot be explained by individual brain areas, but that emerge from the collective coordination between areas at a specific. Synergistic functional interactions therefore reveal the presence of collective behaviour encoding IG that emerge from the coordination of distant brain areas and circuits (Fig. 5B). Contrary to redundant connectivity, synergistic coupling showed a stronger effect across clusters, rather than within clusters (Fig. 6), therefore suggesting a more distributed and long-range nature of synergistic functional interactions. Our results align with a body of literature supporting the idea that cognitive functions, including learning, emerge from the coordinated activity of neural populations distributed across large-scale brain networks (Varela et al., 2001; Bressler and Menon, 2010) and emerge from network-wide and self-organised information routing patterns (Deco et al., 2015; Battaglia and Brovelli, 2020). This distributed nature of neural processing, as observed in task-relevant representations and sensory-to-action transformations, is in line with the growing evidence of brain-wide dynamics during learning and decision-making (Averbeck and Costa, 2017; Bassett and Mattar, 2017; Hunt and Hayden, 2017; Averbeck and O’Doherty, 2021). In mice, recent evidence suggests that transformations linking sensation to action during learning and decision-making are highly distributed and parallelized in a brain-wide manner (Khilkevich et al., 2024).

Within this framework, we suggest that redundant and synergistic functional interactions could be interpreted as a proxy of functional segregation and integration processes, although a direct mapping may be lacking. Redundant functional interactions may appear in collective states dominated by oscillatory synchronisation (Engel, Fries and Singer, 2001; Varela *et al*., 2001; Buzsáki and Draguhn, 2004; Fries, 2015) or resonance phenomena (Vinck *et al*., 2023). Synergistic functional interactions may be associated with functionally-complementary interactions (i.e., functional integration). Indeed, synergistic interactions have been reported between distant transmodal regions during high-level cognition (Luppi *et al*., 2022; Varley, Pope, Faskowitz, *et al*., 2023) and, at the microscale, in populations of neurons within a cortical column of the visual cortex and across areas of the visuomotor network (Nigam, Pojoga and Dragoi, 2019; Varley, Pope, Maria Grazia, *et al*., 2023b). During learning, a recent study has shown that redundant functional interactions encoding either reward or punishment prediction errors are associated by segregated networks in prefrontal regions, whereas the integration between reward and punishment learning is mediated by synergistic interactions between them (Combrisson *et al*., 2023). Synergistic interactions may provide functional advantage, in contrast to redundancy and unique information, because they would enable the full exploitation of possible combinations of neurons or brain areas, making their informational capacity to be exponential with the system size (Rosas *et al*., 2020). Redundancy, instead, would provide robustness, because over-representation would ensure that information remains available even if any neuron or brain region would be perturbed. A drawback of redundancy would be the cost of over-representation, whereas a limitation of synergistic encoding would be the vulnerability to disruptions in individual nodes would disrupt information synergistically held together with other sources (Mediano *et al*., 2022). Indeed, information decomposition analysis of artificial neural networks performing cognitive tasks has shown that while redundant information is required for robustness to perturbations in the learning process, synergistic information is used to combine information from multiple modalities and more generally for flexible learning (Proca *et al*., 2024). Along the same reasoning, artificial networks exhibit higher levels of synergy at comparatively lower orders, rather than higher orders, probably as an attempt to minimise the effect of losing synergistic information with several neurons (Tax, Mediano and Shanahan, 2017). In brain networks, redundant functional interactions may be important for robust sensory and motor functions, whereas synergistic interaction for flexible higher cognition (Luppi *et al*., 2024). The tradeoff between redundancy and synergistic encoding may support robustness and resilience at the large-scale level, as suggested for microscopic neural networks (Panzeri *et al*., 2022). Within this interpretative framework, our results provide supports to the hypothesis that brain interactions regulate segregation and integration processes to support cognitive functions (Sporns, 2013; Braun *et al*., 2015; Deco *et al*., 2015; Cohen and D’Esposito, 2016; Shine *et al*., 2016; Finc *et al*., 2020; Wang *et al*., 2021). Such balanced states may give rise to selective patterns of information flow (Buehlmann and Deco, 2010; Kirst, Timme and Battaglia, 2016; Palmigiano *et al*., 2017; Battaglia and Brovelli, 2020), as observed in the feature-specific information transfer (FIT) results (Figure 8). Taken together, our results suggest general principles for network interactions, whereby synergistic relations encode relevant information for goal-directed learning. The causal effect of synergistic interactions on cognition remains, however, an open question that could be tackled with perturbation or pharmacological manipulations.

### Higher-order synergistic encoding and broadcasting of information gain

In order to explore the nature of synergistic cortico-cortical interactions in directed exploration, we tested whether neuronal interactions supporting cognitive functions require collective behaviours among multiple brain areas, the so-called higher-order correlations (Martignon *et al*., 2000; Schneidman *et al*., 2006; Montani *et al*., 2009; Ohiorhenuan *et al*., 2010; Ganmor, Segev and Schneidman, 2011; Yu *et al*., 2011; Shimazaki *et al*., 2012; Nigam, Pojoga and Dragoi, 2019; Sizemore *et al*., 2019; Panzeri *et al*., 2022). Higher-order interactions are at the core of complex systems (Crutchfield, 1994; Battiston *et al*., 2020, 2021; Santoro *et al*., 2023) and they are thought to support collective behaviours in social systems (Benson, Gleich and Leskovec, 2016), ecology (Grilli *et al*., 2017; Levine *et al*., 2017) and biology (Sanchez-Gorostiaga *et al*., 2019). We generalised information theoretical decomposition methods to higher-order functional interactions, and we showed that information gain is encoded in higher-order synergistic functional interactions up to quadruplets. By definition (Eq. 12), significant multiplets at a given order encode synergistic information that is not present at lower orders. For example, significant triplets contain information beyond what is carried by the underlying pairwise connections. Similarly, the two largest quadruplets—comprising the visual-temporal-vmPFC-OFC and visual-parietal-vmPFC-OFC clusters—encode additional information that cannot be explained by lower-order interactions (Fig. 7A). Our results demonstrate that IG is encoded by functional interactions beyond the pairwise regime. Higher-order synergistic co-modulations encoding IG were found up to order 4. The analysis of centrality of individual brain areas in higher-order behaviours showed that visual areas dominated pairwise functional interactions, whereas the vmPFC-OFC dominated both triplet and quadruplet interactions (Fig. 7C and D). In addition, feature-specific information transfer (FIT) analyses (Celotto *et al*., 2023) showed that IG signals are broadcasted from multiple brain regions and converge to the ventromedial PFC and OFC (Fig. 8). These results further support the conclusion highlighting a central role of the vmPFC-OFC cluster in the processing of information gain.

### Limitations and open questions

Information gain leads to uncertainty reduction in action-outcome associations. People are equipped with cognitive systems that estimate and handle uncertainty (Bach and Dolan, 2012) to drive further learning (Daw, Niv and Dayan, 2005; Yoshida and Ishii, 2006; Behrens *et al*., 2007; Payzan-LeNestour and Bossaerts, 2011; Payzan-LeNestour *et al*., 2013). Uncertainty can be formalised as the entropy of the probability distribution over possible actions, representing the degree of unpredictability about most rewarding action. From the mathematical point of view, however, uncertainty and information gain are related. Information gain can be decomposed into the difference of the cross entropy of *P*(*a*_*t*−1_) and *P*(*a*_*t*_), which measures the mismatch between the prior and updated posterior policy, and the entropy of the posterior distribution *P*(*a*_*t*_), which measures the remaining uncertainty in the action selection policy after update. The difference of these terms measures the information gain or the reduction in uncertainty achieved by updating the prior to the posterior after each observation (see methods section for more details). IG and uncertainty reduction are closely linked, particularly in the deterministic task used in this study. Further research is needed to explore whether IG and uncertainty signals are differently encoded in synergistic interactions in the brain.

A second limitation is the difficulty in determining whether information gained from correct outcomes differs from that gained from errors. In this task, IG and RPEs show a U-shaped relationship, with both errors (negative RPE) and successes (positive RPE) linked to positive IG. As learning progresses, both RPE and IG tend to decrease. Therefore, we cannot determine if IG is more associated with confirmation (linked to positive RPEs) or conflict monitoring (linked to negative RPEs). The IG signals here reflect a combined contribution from both errors and successes. Future research could explore the distinct circuits involved in IG from errors and successes during goal-directed learning.

A third limitation is the focus on high-gamma activity, as it may overlook the role of phase synchronization in neural interactions. Phase synchronization, particularly in lower frequency bands (such as alpha and beta range), facilitates long-range connectivity by aligning neural excitability across regions, enabling efficient information transfer (Engel et al., 2001; Varela et al., 2001; Brovelli et al., 2004; Buzsáki and Draguhn, 2004; Fries, 2015; Vinck et al., 2023). Ignoring these lower-frequency phase interactions may lead to an incomplete understanding of large-scale network dynamics and neural coordination supporting cognitive processes. Further studies are needed to bridge the explanatory gap between neural interactions supported by high-gamma activity and phase-related synchronization in brain networks.

An open question concerns the role of the basal ganglia and cortico-striatal interactions in the encoding of information gain. The inherent limitations of MEG in resolving deep brain structures prevent a detailed study of these interactions. Intracranial EEG recordings in epileptic patients rarely include electrodes in the striatum, the main input structure of the basal ganglia. As a result, human neurophysiology provides limited access to fast neural dynamics and cortico-striatal interactions. The absence of measurements in the basal ganglia could explain why we did not observe correlates of reward prediction error (RPE), as this region plays a crucial role in encoding reward signals. Future studies investigating the functional role of cortico-striatal interactions in information seeking and goal-directed learning in primates will be crucial for advancing our understanding of adaptive behavior and decision-making processes.

Although the number of participants (N=11) is limited, our previous study indicated that it has appropriate group-level statistical power (Combrisson et al., Neuroimage, 2019). In the study, the Matthews Correlation Coefficient (MCC) was used to determine the minimum number of trials and subjects for group-level inference with cluster-wise corrections. An MCC of 0.8 or higher indicates excellent model performance. With an average of 225 trials per participant, the MCC was approximately 0.9. Nevertheless, we cannot exclude that significant correlates of RPE could be identified if a larger number of participants could be studied.

Finally, the Q-learning model was used to estimate the trial-by-trial variability in RPE and IG signals, rather than to serve as a comprehensive model of choice behaviours in our task. Indeed, the model does not account for the win-stay and lose-switch strategies observed in our task. As standard RL models rarely capture across-trial behavioral strategies, accounting for it would require new computational theories whose development is beyond the scope of this work.

To conclude, the current study provides evidence that information gain is encoded in synergistic and higher-order functional interactions alongside with its broadcasting towards the prefrontal reward circuitry. Our study provides evidence regarding how information relevant for cognition is encoded and broadcasted in distributed cortical networks and brain-wide dynamics.

## METHODS

### Experimental conditions and behavioural learning task

Eleven healthy subjects participated in the study (all right handed, 7 females; average age 26 years old). All participants gave written informed consent according to established institutional guidelines, and they received monetary compensation (50 euros) for their participation. The project has been promoted by the INSB of the CNRS (CNRS No. 17020; ANSM No. 2017-A03614-49) and approved by the ethical committee CPP Sud-Méditerranée.

Goal-directed learning was studied using an arbitrary visuomotor learning task design, where the relation between the visual stimulus, the action and its outcome is arbitrary and causal. Participants were instructed to learn by trial-and-error the correct associations between 3 coloured circles and 5 finger movements (Fig. 1A). Participants were instructed that correct actions were not exclusive across stimuli, meaning that they could not exclude certain fingers if assigned to another stimulus when seeking the correct association. On each trial, participants were presented a coloured circle to which they had to respond within 1.5 s. Reaction times were computed as the time difference between stimulus presentation and motor response (finger movement). After a fixed delay of 1s following the disappearance of the colored image, an outcome image was presented for 1s and informed the subject whether the response was correct, incorrect, or too late (if the reaction time exceeded 1.5s). “Late” trials were excluded from the analysis, because they were either absent or very rare (i.e., maximum 2 late trials per session). The next trial started after a variable delay ranging from 2 to 3 s (randomly drawn from a uniform distribution) with the presentation of another visual stimulus. Visual stimuli (colored circles) were randomised in blocks of three trials. Each learning session was composed of 60 trials, 3 stimulus types (i.e., different colours, S1, S2, and S3) and 5 possible finger movements. Each subject performed 4 learning sessions, each containing different sets of coloured stimuli. To ensure reproducible performances across sessions and subjects, the task design manipulated learning and produced reproducible phases of acquisition and consolidation across sessions and individuals (Brovelli *et al*., 2008). More precisely, the correct stimulus–response associations were not set a priori. Instead, the correct stimulus–response associations were assigned as the subject proceeded in the task. Figure 1B shows the matrix of all the possible stimulus–response combinations, updated according to the exemplar learning session (Fig. 1A). The first presentation of each stimulus was always followed by an incorrect outcome, irrespective of the subject’s motor response (Fig. 1B, from trials 1 to 3). Then, on the second presentation of stimulus S1, any untried finger movement was always considered as a correct response. Because the stimuli were presented randomly within blocks of 3 trials, stimulus S1 could occur for the second time from trials 4 to 6 (trial 4 in Fig. 1). For the second stimulus S2, the response was correct only when the subjects had performed 3 incorrect finger movements (trial 9 in Fig. 1B). For stimulus S3, the subject had to try 4 different finger movements before the correct response was found (trial 12 in Fig. 1B). In other words, the correct response was the second finger movement (different from the first tried response) for stimulus S1, the fourth finger movement for stimulus S2, the fifth for stimulus S3. Each learning session was composed of 42 trials, 3 stimulus types (i.e., different colors, S1, S2, and S3) and 5 possible finger movements. During scanning, each subject performed 4 learning sessions, each containing new colored stimuli.

### MarsAtlas cortical parcellation and source model

Anatomical T1-weighted MRI images were acquired for all participants using a 3-T whole-body imager equipped with a circular polarised head coil (Bruker). Magnetoencephalographic (MEG) recordings were performed using a 248 magnetometers system (4D Neuroimaging magnes 3600). Visual stimuli were projected using a video projection and motor responses were acquired using a LUMItouch® optical response keypad with five keys. Presentation® software was used for stimulus delivery and experimental control during MEG acquisition.

Single-subject cortical parcellation was performed using the MarsAtlas approach (Auzias, Coulon and Brovelli, 2016). After denoising using non-local means (Coupe *et al*., 2008), T1-weighted MR-images were segmented using the FreeSurfer “recon-all” pipeline (v5.3.0). Grey and white matter segmentations were imported into the BrainVisa software (v4.5) and processed using the Morphologist pipeline procedure (http://brainvisa.info). White matter and pial surfaces were reconstructed and triangulated, and all sulci were detected and labelled automatically (Mangin et al., 2004). A parameterization of each hemisphere white matter mesh was performed using the Cortical Surface toolbox (http://www.meca-brain.org/softwares/). It resulted in a 2D orthogonal system defined on the white matter mesh, constrained by a set of primary and secondary sulci (Auzias et al., 2016). The MarsAtlas parcelization contained a total of 82 cortical parcels (41 each hemisphere). Each individual white matter mesh was then resampled onto a template defined by the average geometry across a population of 138 healthy subjects (https://meca-brain.org/software/hiphop138/) in two steps: 1) the parameterization induced by the 2D orthogonal system is used to map the white matter mesh onto a canonical spherical domain; 2) the location of each vertex of the template mesh is interpolated onto the individual mesh geometry using the classical barycentric mapping. This procedure defines one-to-one spatial correspondence across the white matter meshes from all individuals at the vertex level. By adapting the spatial resolution of the template mesh, we can fix the number of points that define the correspondence across subjects. The sources are then defined at the location of these points and are thus distributed at consistent anatomical locations across subjects.

Finally, we created source space and forward models readable by MNE-python (Gramfort *et al*., 2013) using cortical meshes generated using BrainVISA (MarsAtlas) and subcortical structures generated by Freesurfer. These three elements are needed for the power estimation at the source level, which will be discussed in the next paragraph. These steps were performed using the BV2MNE toolbox (https://github.com/brainets/bv2mne), a python library developed in our team based on MNE-python . The anatomical meshes were coregistred with the MEG data using the ‘mne coreg’ interface.

### Single-trial high-gamma activity (HGA) in MarsAtlas

We focused on broadband gamma from 60 to 120Hz for two main reasons. First, it has been shown that the gamma band activity correlates with both spiking activity and the BOLD fMRI signals (Mukamel *et al*., 2005; Niessing *et al*., 2005; Lachaux *et al*., 2007; Nir *et al*., 2007; Ray and Maunsell, 2011), and it is commonly used in MEG and iEEG studies to map task-related brain regions (Brovelli *et al*., 2005; Crone, Sinai and Korzeniewska, 2006; Jerbi *et al*., 2009). Therefore, focusing on the gamma band facilitates linking our results with the fMRI and spiking literature on probabilistic learning. Second, single-trial and time-resolved high-gamma activity can be exploited for the analysis of cortico-cortical interactions in humans using MEG and iEEG techniques (Brovelli *et al*., 2015, 2017; Combrisson, Allegra, *et al*., 2022; Combrisson *et al*., 2024).

MEG signals were high pass filtered to 1Hz, low-pass filtered to 250 Hz, notch filtered in multiples of 50Hz and segmented into epochs aligned on stimulus and outcome presentation. Independent Component Analysis (ICA) was performed to detect and reject cardiac, eye-blink and oculomotor artefacts. Artefact rejection was performed semi automatically, at first by a visual inspection of the epochs’ time series, then by means of the *autoreject* python library (Jas *et al*., 2017) to detect and reject bad epochs from further analysis.

Spectral density estimation was performed using a multi-taper method based on discrete prolate spheroidal (slepian) sequences (Percival and Walden, 1993). To extract high-gamma activity from 60 to 120Hz, MEG time series were multiplied by *k* orthogonal tapers (*k* = 11) (0.2s in duration and 60Hz of frequency resolution, each stepped every 0.005s), centred at 90Hz and Fourier-transformed. Complex-valued estimates of spectral Measures 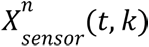, including cross-spectral density matrices, were computed at the sensor level for each trial *n*, time *t* and taper *k*. Source analysis requires a physical forward model or leadfield, which describes the electromagnetic relation between sources and MEG sensors. The leadfield combines the geometrical relation of sources (dipoles) and sensors with a model of the conductive medium (i.e., the head model). For each participant, we generated a boundary element model using a single-shell model constructed from the segmentation of the cortical tissue obtained from individual MRI scans (Nolte, 2003). Lead-fields were not normalised. The head model, source locations and the information about MEG sensor position for both models were combined to derive single-participant leadfields. The orientation of cortical sources was set perpendicular to the cortical surface.

We used adaptive linear spatial filtering to estimate the power at the source level. In particular, we employed the Dynamical Imaging of Coherent Sources (DICS) method, a beamforming algorithm for the tomographic mapping in the frequency domain (Gross *et al*., 2001), which is a well suited for the study of neural oscillatory responses based on single-trial source estimates of band-limited MEG signals. At each source location, DICS employs a spatial filter that passes activity from this location with unit gain while maximally suppressing any other activity. The spatial filters were computed on all trials for each time point and session, and then applied to single-trial MEG data.

Single-trial power estimates aligned with outcome and stimulus onset were log-transformed to make the data approximate Gaussian and low-pass filtered at 50Hz to reduce noise. Single-trial mean power and standard deviation in a time window from -0.5 and -0.1 s prior to stimulus onset was computed for each source and trial, and used to z-transform single-trial movement-locked power time courses. Similarly, single-trial outcome-locked power time courses were log-transformed and z-scored with respect to baseline period, to produce HGAs for the prestimulus period from -1.6 to -0.1 s with respect to stimulation for subsequent functional connectivity analysis. Finally, single-trial HGA for each brain region of *MarsAtlas* was computed as the mean z-transformed power values averaged across all sources within the same region. Single-trial HGA estimates were performed using home-made scripts based on MNE-python (Gramfort *et al*., 2013).

### Model-based analysis of behavioural data and estimate of learning signals

We estimated the evolution of stimulus-action-outcome probabilities and updating learning signals during learning using a Q-learning model (Watkins and Dayan, 1992) from reinforcement learning theory (Sutton and Barto, 2018). Briefly, the Q-learning model updates action values through the Rescorla-Wagner learning rule (1972) expressed by the following equation:

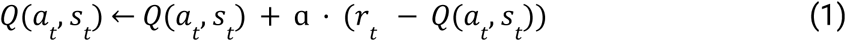

The left-hand side of the equation corresponds to the updated Q-value for the observed stimulus *s*_*t*_ and chosen action *a*_*t*_ after the observation of the outcome *r*_*t*_, at trial *t*. In our case, *s*_*t*_ corresponds to one of the three coloured stimuli, while *a*_*t*_ represents one of the five possible finger movements and *r*_*t*_ is either one or zero for correct and incorrect outcomes, respectively. Q-values were then transformed into probabilities according to the softmax equation :

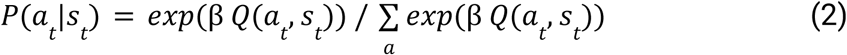

*P*(*a*_*t*_ |*s*_*t*_) is the conditional probability of selecting action *a* given stimulus *s*, and it sums to one over possible actions 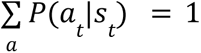. For simplicity, we drop the conditioning over stimulus *s* and refer to it as action policies *P*(*a*_*t*_). The coefficient β is termed the inverse ‘temperature’: low β (less than 1) causes all actions to be (nearly) equiprobable, whereas high β (greater than 1) amplifies the differences in association values. We identified the set of parameters that best fitted the behavioural data using a maximum likelihood approach, as in the previous paper (Brovelli et al., 2008). The model was fitted separately for each block of trials and learning session. For each learning session, we used a grid-search algorithm varying the learning rate a from 0.1 to 1 (in steps of 0.01) and the inverse temperature β from 1 to 10 (in steps of 0.2). Generally, grid-search algorithms are often preferred over optimization methods for parameter inference, because they systematically explore the entire parameter space, ensuring that the best possible combination is found. Unlike optimization techniques, which may get stuck in local minima, grid search guarantees a comprehensive evaluation of all parameter choices. This exhaustive search is particularly useful when dealing with low-dimensional problems or when prior knowledge about the best region of the parameter space is limited. In addition, a previous fMRI study used the same task and Q-learning model (Brovelli et al., 2008), and it showed that the grid-search approach provided accurate estimates of action-outcome conditional probabilities that fitted behavioral data (Figure 2 and Figure 3 in Brovelli et al 2008). The optimal set of parameters was chosen so to maximise the log-likelihood of the probability to make the action performed by the participant *P*(*c*_*t*_) = *P*(*a*_*t*_ = *chosen*), defined as 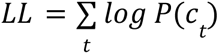. Model fitting was performed per learning block, so to accommodate potential changes in ɑ and β across sessions. Therefore, the Q-learning model provides estimates of the evolution of reward prediction-errors during learning as:

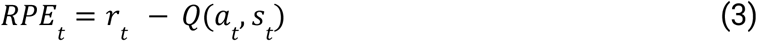

Bayesian inference is a principled statistical model based on Bayesian updating of beliefs about the current state of the world. The amount of update after each observable is related to the notion of information gain. In Bayesian statistics, information gain can be defined as the amount of information required to “revise” one’s beliefs from the prior probability distribution *P*(*a*_*t*−1_) to the posterior probability distribution *P*(*a*_*t*_). This measure can be formalised via the Kullback–Leibler (KL) divergence between the posterior and prior. In neuroscience, this measure is also referred to as the Bayesian surprise (Itti and Baldi, 2009; Modirshanechi *et al*., 2023):

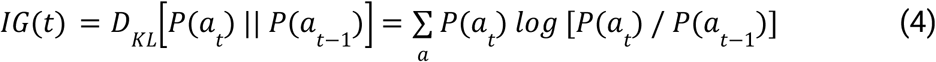

Note that the information gain depends on the stimulus presented at each trial. The index associated with stimulus type was dropped for simplicity. The equation measuring information gain can be expanded and rewritten as

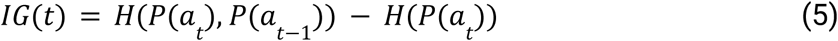

The first term on the right-hand side is the cross entropy of *P*(*a*_*t*−1_) and *P*(*a_t_*), and it measures the mismatch between the prior and updated posterior policy. The second term is the entropy of the posterior distribution *P*(*a_t_*) and it measures the remaining uncertainty in the action selection policy after update. The difference of these terms measures the information gain or the reduction in uncertainty achieved by updating the prior to the posterior after each observation. Information gain therefore relates to epistemic uncertainty, as it captures the reduction of model uncertainty due to observed outcomes. In the current task, information gain and uncertainty are correlated, because the correct action policies are deterministic. During the initial exploratory phase, both information gain and uncertainty start high when the participant’s policy is far from optimal and uncertain. At the end of learning, the policy becomes approximately deterministic, and both information gain and uncertainty approach zero. Information gain after the observation of the outcome leads to a reduction of uncertainty in the policy. No dissociation can be drawn between IG and uncertainty reduction in the current task. Nevertheless, information gain and reward prediction errors are not linearly related. Errors (i.e., negative RPE) and successes (i.e., positive RPE) are both associated with positive IG. In the current task, IG and RPE display approximately a U-shape relationship. On the other hand, IG scales with the absolute value of RPE, because both reflect the impact of surprise on learning. The absolute value of RPE captures how unexpected an outcome is. IG quantifies how much uncertainty is reduced as a result of that unexpected outcome.

### Cortical areas encoding learning signals

We used information-theoretic metrics to quantify the statistical dependency between single-trial HGA and the outcome-related learning signals. Information-based measures quantify how much the neural activity of a single brain region explains a variable of the task. To this end, we computed the mutual information (MI); as a reminder, mutual information is defined as:

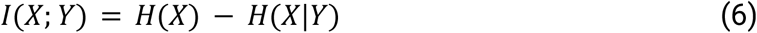

In this equation the variables *X* and *Y* represent the HGA power and the behavioural variables, respectively. *X* represents the HGA power over time, trials and regions. *Y* represents the behavioural variables across trials. *H*(*X*) is the entropy of *X*, and *H*(*X*|*Y*) is the conditional entropy of *X* given *Y*. We used Gaussian-Copula Mutual Information (GCMI) (Ince *et al*., 2017), which is a semi-parametric binning-free technique to calculate MI. In the GCMI approach, which includes a rank based normalisation of the data, entropy values are computed using the differential entropy formula for a Gaussian distribution. The transformed data allows entropy estimation based on the determinant of the covariance matrix and the assumption of Gaussian marginals. The GCMI is a robust rank-based approach that allows detecting any type of monotonic relation between variables. More precisely, the GCMI is a lower-bound approximate estimator of mutual information for continuous signals based on a Gaussian copula, and it is of practical importance for the analysis of short and potentially multivariate neural signals. Note, however, that the GCMI does not detect non-monotonic (e.g., parabolic) relations. In the current work, the GCMI was computed across trials between time-resolved HGA and outcome-related learning signals. The parametric GCMI estimation was bias-corrected using an analytic correction to compensate for the bias due to the estimation of the covariance matrix from limited data (i.e. here, limited number of repetitions or trials) (Ince et al., 2017). Since this parametric correction only depends on the number of trials, the same value is going to be used for both permuted and non-permuted data. Therefore, this bias correction only impacts the estimated effect size but has no effects on statistical results. The considered outcome-related learning signals were reward prediction errors (RPE) and Information Gain (IG).

### Cortico-cortical functional interactions encoding learning signals

The goal of network-level analyses was to identify dynamic functional networks encoding outcome-related computations, such as the reward prediction errors (RPE) and information gain (IG). We used information-theoretic metrics to quantify the statistical dependency between single-trial event-related co-modulations in HGA across trials and learning computations. In general terms, the aim of our analyses was to assess the interdependence between pairs of brain signals (X_1_ and X_2_) and a third learning variable (Y), either RPE or IG. More precisely, we wished to quantify whether the interaction in activity co-modulation between brain regions supported functional segregation and integration phenomena supporting learning. We reasoned that functional segregation could be quantified as the amount of redundant information shared in the co-modulation of activity across brain regions. Similarly, we propose that functional integration processes could be quantified as the amount of synergistic information carried by the two brain regions about the target learning variable. Such research questions can be formalised within the Partial Information Decomposition (PID) framework (Williams and Beer, 2010; Lizier *et al*., 2018). Indeed, the PID framework allows the decomposition of multivariate mutual information between a system of predictors and a target variable, and to quantify the information that several sources (or predictors) variables provide *uniquely*, *redundantly* or *synergistically* about a target variable. In the current study, we considered the case where the source variables are the across-trials HGA of two brain regions (X_1_ and X_2_) and the target variable is the trial-by-trial evolution of the learning variable (Y). If pairs of brain regions X_1_ and X_2_ carry the same information about learning variable Y, we refer to it as *redundant* information. If only one area X_1_ or X_2_ carries information about Y, it is referred to as *unique* information carried by X_1_ or X_2_ about Y, and vice-versa for Y. Finally, if neither of the variables alone provide information about Y, but they need to be observed together, we refer to it as *synergistic* information. In other words, the knowledge of any of the predictors separately does not provide any information about the target variable C. Analytically, the PID proposes to decompose the total mutual information between a pair of source variables X and Y and a target variable Z *I*(*X*_1_, *X*_2_; *Y*) into four non-negative components:

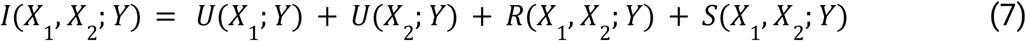

*U*(*X*_1_;*Y*) and *U*(*X*_2_;*Y*) are unique information carried for the two areas, respectively. *R*(*X*_1_,*X*_2_;*Y*)*S*(*X*_1_,*X*_2_;*Y*) are the redundancy and synergy terms, respectively. In addition, the PID formulation links to classical Shannon measures of mutual information as

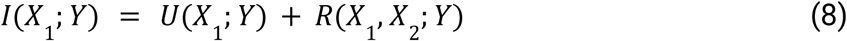

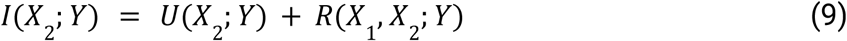

The problem with this approach is that its governing equations form an under-determined system: only three quantities that can be computed from the data (i.e., the mutual information quantities *I*(*X*_1_, *X*_2_; *Y*), *I*(*X*_1_; *Y*) and *I*(*X*_2_; *Y*)) for the four terms of the PID (i.e., two unique information, redundant and synergistic). To actually calculate the decomposition, an additional assumption must therefore be made. Here, we exploit the so-called minimum mutual information (MMI) PID, which has been shown to provide correct estimations for a broad class of systems following a multivariate Gaussian distribution (Barrett, 2015; Luppi *et al*., 2022). According to the MMI PID, redundant information carried by pairs of brain regions is given by the minimum of the information provided by each individual source to the target,

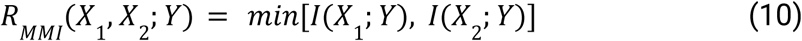

Then, synergistic information can be computed by substituting Eq. 8, 9 and 10 in Eq. 7 and rearranging the terms

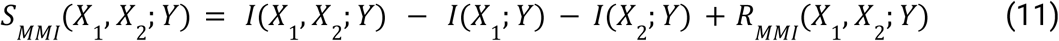

Equations 9 and 10 represent the redundant and synergistic information carried by the co-modulations in HGA of pairs of brain regions X_1_ and X_2_ about the learning variable Z, respectively. By definition, redundant functional interactions carry the same information about learning variables than the one carried by individual brain areas. On the other hand, synergistic connectivity reveals brain functional interactions that cannot be explained by individual contribution, but only by their combined and collective coordination.

### Higher-order cortico-cortical interactions encoding learning signals

We then studied whether outcome-related learning signals would be encoded in the interactions beyond the pairwise relations, the so-called higher-order functional interactions, or correlations. The definitions of redundancy and synergy can be generalised to higher-order by simply iterating over all regions of a multiplet. By considering a multiplet as a set of *n* variables *X*^*n*^ = {*X*_1_, …, *X_n_*}, the redundant information carried by each higher-order multiplet about the target variable is given by:

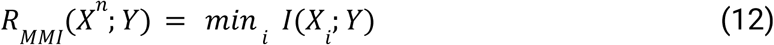

Then, synergistic information can be computed by substituting Eq. 8, 9 and 10 in Eq. 7 and rearranging the terms

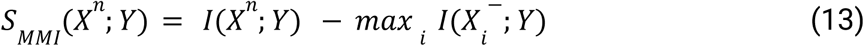

where *X_i_*^−^ is the the set of variables of *X^n^* excluding the brain area *i*. One of the main issues in the study of the higher-order behaviours of complex systems is the computational cost required to investigate all the multiplets of any order. In order to perform HOIs analyses over all orders, we thus identified 9 sets of brain regions that displayed significant encoding of learning signals and we averaged the HGA for each set of regions-of-interest. This allowed us to perform an exhaustive analysis of HOIs at all orders. Equations 11 and 12 represent the redundant and synergistic information carried by multiplets of brain regions about the learning variable Y, respectively.

### Cortico-cortical information transfer encoding learning signals

In order to investigate whether learning signals are broadcasted across different cortical regions, we used a recently-developed information-theoretic measure termed Feature-specific Information Transfer (FIT) that quantifies how much information about specific features flows between brain areas (Celotto *et al*., 2023). FIT merges the Wiener-Granger causality principle (Granger, 1980; Brovelli *et al*., 2004; Bressler and Seth, 2011) with content specificity based on the PID framework (Williams and Beer, 2010; Lizier *et al*., 2018). FIT isolates information about a specific task variable Y encoded in the current activity of a receiving neural population, which was not encoded in its past activity, and which was instead encoded by the past activity of the sender neural population. More precisely, the FIT measure is based on two four-variables PID and it is defined as the minimum of two “atoms”. The first atom is defined in a four-variable PID, which considers the learning variable *Y* as the target and three source variables, namely the brain activity in the past of the sending signal *X*_1_ (*t* − *τ*), the past of the receiver *X*_2_ (*t* − *τ*) and the present of the receiver *X*_2_ (*t*). This atom formalises the principle of Granger-Wiener causality within the PID framework and it adds the encoding of task variables (e.g., learning) in the time-lagged predictability between signals. More formally, the first atom is defined as the redundant information about *Y* carried by the past of *X*_1_ (*t* − *τ*) and the present of *X*_2_ (*t*), which is unique with respect to past of the *X*_2_ (*t* − *τ*). As for previous analyses, we used the MMI PID to compute this atom, which was computed as

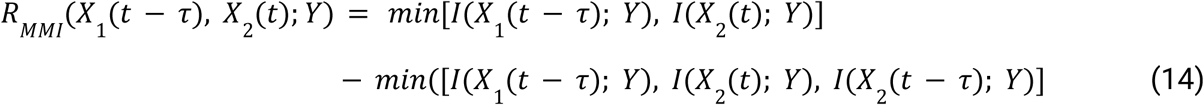

Note that the second term on the right-hand side assures the uniqueness with respect to the past of the receiver *X*_2_ (*t* − *τ*). The second atom for the calculation of the FIT is defined for a four-variable PID, considering the present of the receiver *X*_2_ (*t*), as the target variable and the learning variable *Y*, *X*_1_ (*t* − *τ*) and *X*_2_ (*t* − *τ*) as source variables. This second term assures that the FIT measure does exceed: i) the overall propagation of information between signals, referred to as the Directed Information (Massey 1990) or Transfer Entropy (Schreiber, 2000), and quantified by means of the conditional mutual information *I*(*X*_1_ (*t* − *τ*); *X*_2_ (*t*)|*X*_2_ (*t* − *τ*)); ii) the mutual information between the learning variable and the past of the sender, *I*(*Y*; *X*_1_ (*t* − *τ*)) and iii) the mutual information between the learning variable and the present of the receiver, *I*(*Y*; *X*_2_ (*t*)). This term is defined as the redundant information about the present of the receiver *X*_2_ (*t*) carried by *X*_1_ (*t* − *τ*) and *Y*, which is unique with respect to the past of the receiver *X*_2_ (*t* − *τ*):

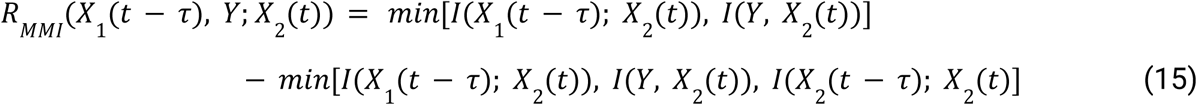

Therefore, the FIT measure is defined as the minimum between these two atoms (Eq. 14 and 15) to select the smallest nonnegative piece of information:

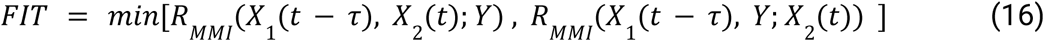

We computed the FIT measure between all pairs of 8 sets of brain regions at different delays tau ranging from -0.005 to -0.2s, and averaged over delays. Given that the scope of the study was not a detailed characterisation of time delays between clusters, we used such averaging procedure, as in previous papers (Lemke et al, 2024; Celotto et al, 2023). Statistical analysis was similarly applied as for all other measures.

### Statistical analysis

For the statistical inferences, we used a group-level approach based on non-parametric permutations and encompassing non-negative measures of information developed in a previous study (Combrisson, Allegra, *et al*., 2022). We used a random effect (RFX) to take into account the inter-subject variability. In this approach, the information theoretical metrics (e.g., mutual information, MI) between the neurophysiological signal and the behavioural regressor are computed across trials for each participant separately, at each time point and brain region. In order to assess whether the estimated effect size in MI significantly differed from the chance distribution and to correct for multiple comparisons, we implemented a cluster-wise statistics approach (Maris and Oostenveld, 2007). For cluster-level statistics, the cluster forming threshold is defined as the 95th percentile across all of the permutations. Such threshold is used to identify the clusters on both the true and permuted data. To sample the distribution of MI attainable by chance, we computed the MI between the brain data and a randomly shuffled version of the behavioural variable. This procedure was then repeated 1000 times. We took the mean of the MI values computed on the permutations, and used this mean (MI) to perform a one sample t-test across all the participants’ MI values obtained both from original and permuted data. We then used a cluster-based approach to assess whether the size of the estimated t-values significantly differs from its distribution. The cluster forming threshold was defined as the 95th percentile of the distribution of t-values. We used this cluster forming threshold to identify the cluster mass of t-values on both original and permuted data. As a reminder, the cluster-based approach (Maris and Oostenveld, 2007) is performed on the mass of the clusters (i.e., the summed activity above the threshold), rather than on individual temporal samples. Thus, the p-value of the cluster is not representative of the individual samples within the cluster and cannot be interpreted as a measure of the onset and offset of significant effects. Finally, to correct for multiple comparisons across both time and space, we build a distribution made of the 1000 largest clusters estimated on the permuted data. The final corrected p-values were inferred as the proportion of permutations exceeding the t-values. For what concerns functional connectivity analysis, the information theoretical metrics of interest (e.g., redundant connectivity) across pairs of brain regions were studied similarly as the local MI time courses.

## Data availability

The group-level data and results, in addition to Jupiter notebooks to reproduce the figures of the paper, have been deposited on the Github repository (https://github.com/brainets/hosi_infogain) (see Brovelli et al (2025). https://doi.org/10.5281/zenodo.15674302). The single-subject MRI, MEG and behavioral data are protected and are not available due to data privacy laws. The processed data may be requested to the corresponding author.

## Code availability

Higher-order redundancy and synergistic measures were computed using the HOI toolbox (https://github.com/brainets/hoi). All pair-wise information theoretical measures, including the FIT, and statistical analyses were performed using the Frites Python package (https://github.com/brainets/frites) (Combrisson, Basanisi, *et al*., 2022). Hypergraph visualisation was performed using the XGI package (https://xgi.readthedocs.io/en/stable/).

## Acknowledgements

A.B. and E.C were supported by the PRC project “CausaL” (ANR-18-CE28-0016) and received funding from the European Union’s Horizon 2020 Framework Programme for Research and Innovation under the Specific Grant Agreement No. 945539 (Human Brain Project SGA3). A.B. was supported by A*Midex Foundation of Aix-Marseille University project “Hinteract” (AMX-22-RE-AB-071). R.B. and M.N. have received funding from the French government under the “France 2030” investment plan managed by the French National Research Agency (reference : ANR-16-CONV000X / ANR-17-EURE-0029) and from Excellence Initiative of AixMarseille University - A*MIDEX (AMX-19-IET-004). The “Center de Calcul Intensif of the Aix-Marseille University (CCIAM)” is acknowledged for high-performance computing resources. A.B, E.C and D.M. were supported by EU’s Horizon 2020 Framework Programme for Research and Innovation under the Specific Grant Agreements No. 101147319 (EBRAINS 2.0 Project). G.A. was supported by the ANR SulcalGRIDS Project (ANR-19-CE45-0014). We thank Pedro Mediano and Luca Faes for fruitful suggestions and discussions.

## Author contribution statements

Conceptualization: AB

Data curation: AB, EC, RB, MN, GA

Formal analysis: AB, RB

Funding acquisition : AB, RB, GP, DM, SP

Investigation: AB, RB

Methodology: All authors

Project administration: AB

Resources: AB, SP

Software: EC, RB, GA, MN

Supervision: AB, DM, GP

Visualization: AB, EC, RB

Writing (original draft): AB

Writing (review & editing): All authors

