## Supplementary material for "Higher-order and distributed synergistic functional interactions encode information gain in goal-directed learning": Suppl Material

### Supplementary figures

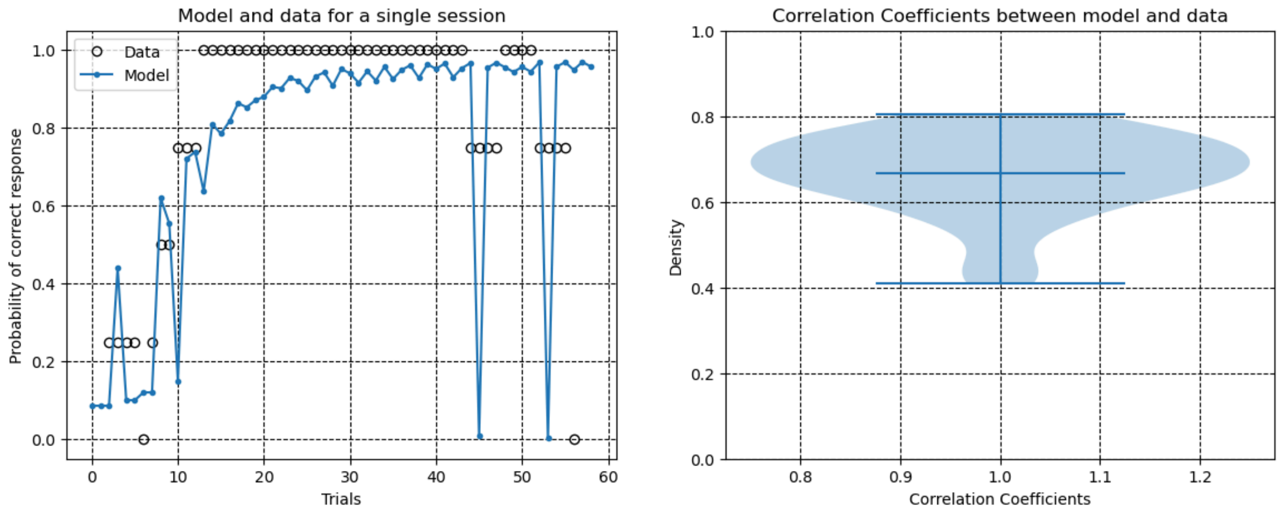


**Supplementary Figure 1. Behavioural results and model comparison.** (A) Comparison between the probability of correct response predicted by the model $P(a_t|s_t)$ as described in Eq. 2 (blue curve) and its data-driven equivalent computed (empty circles), computed as a moving average of the sequence of binary outcomes. The panel shows an exemplar session and it shows how the model (blue curve) is able to track behavioural data (black circles). (B) Violin plot of the distribution across sessions (Ns=4) and participants (N=11) of Pearson's correlation coefficient values between the probability of correct response computed on behavioural data and predicted by the model. The minimum value of correlation coefficient was 0.41 (p=0.003).

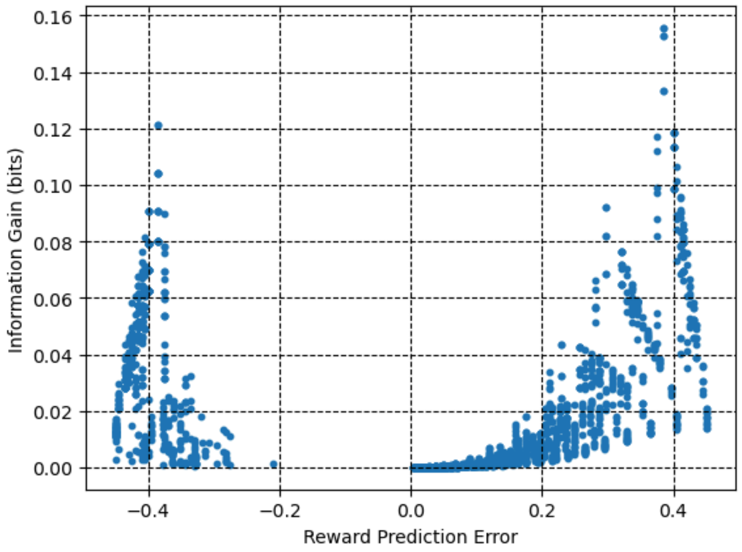


**Supplementary Figure 2. Relation between Information Gain (y-axis) and Reward Prediction Error (RPE, x-axis) signals.** IG and RPE display a U-shape relationship. Errors (i.e., negative RPE) and successes (i.e., positive RPE) are both associated with positive IG. As learning advances, both RPE and IG tend to zero.

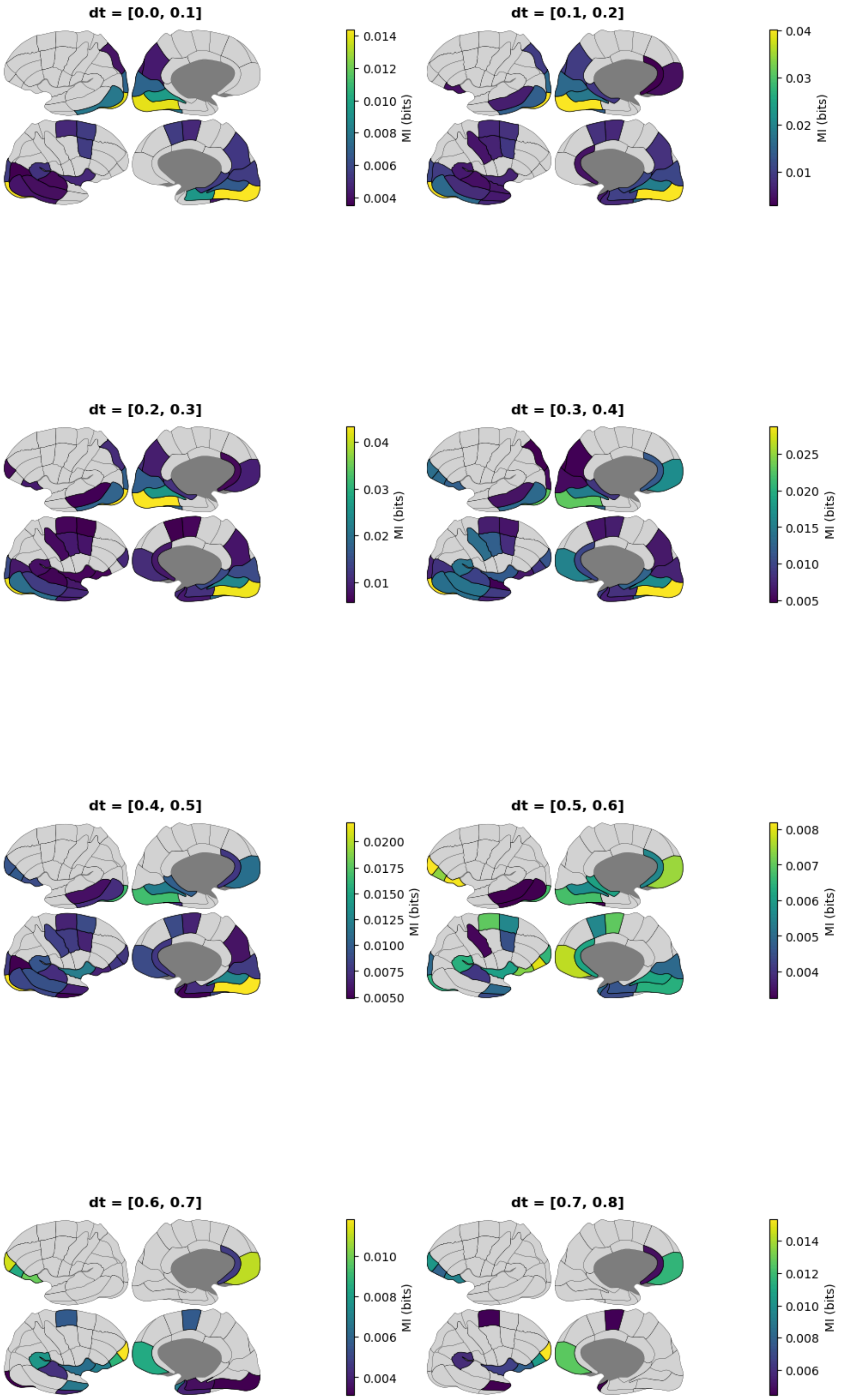


**Supplementary Figure 3. Group-level statistics and local spatio-temporal correlates of information gain (IG).** Regions-of-interest (ROI) of the MarsAtlas (Auzias et al., 2017) showing a significant relation with IG are depicted on an inflated brain plot. In each panel, colored areas indicate the average mutual information between HGA and IG. Shaded grey areas are not significant. The title of each panel represents the time interval considered.

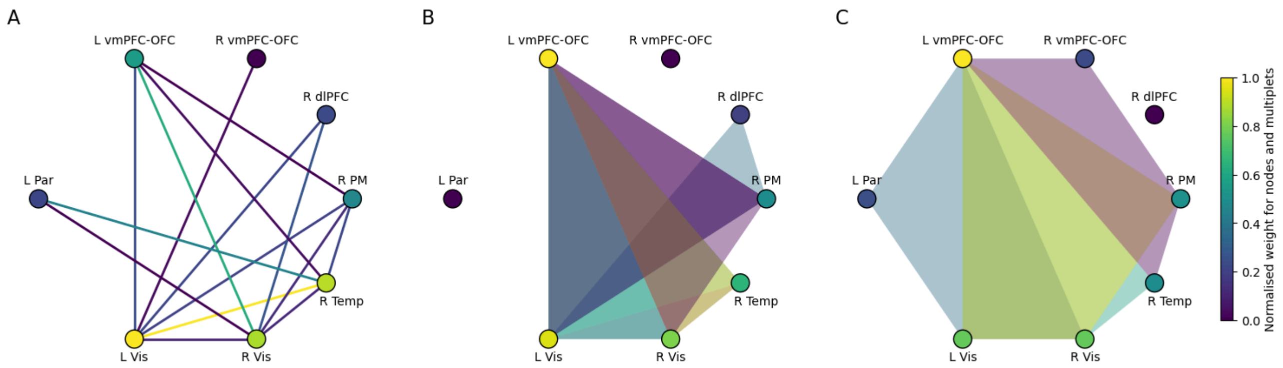


**Supplementary Figure 4. Synergistic Encoding of IG by Brain Networks and Higher-Order Interactions.** Each panel displays eight clusters, with node color representing the normalized average weight of each node. (A) Pairwise synergistic interactions encoding IG. (B) Triplet interactions and (C) quadruplet interactions, visualized as hypergraphs with shaded areas proportional to the average weight.
